# The genome of a daddy-long-legs (Opiliones) illuminates the evolution of arachnid appendages and chelicerate genome architecture

**DOI:** 10.1101/2021.01.11.426205

**Authors:** Guilherme Gainett, Vanessa L. González, Jesús A. Ballesteros, Emily V. W. Setton, Caitlin M. Baker, Leonardo Barolo Gargiulo, Carlos E. Santibáñez-López, Jonathan A. Coddington, Prashant P. Sharma

**Author notes:** co-first authors; corresponding authors, **Email:**.

## Abstract

Chelicerates exhibit dynamic evolution of genome architecture, with multiple whole genome duplication events affecting groups like spiders, scorpions, and horseshoe crabs. Yet, genomes remain unavailable for several chelicerate orders, such as Opiliones (harvestmen), which has hindered comparative genomics and developmental genetics across arachnids. We assembled a draft genome of the daddy-long-legs *Phalangium opilio,* which revealed no signal of whole genome duplication. To test the hypothesis that single-copy Hox genes of the harvestman exhibit broader functions than subfunctionalized spider paralogs, we performed RNA interference against *Deformed* in *P. opilio.* Knockdown of *Deformed* incurred homeotic transformation of the two anterior pairs of walking legs into pedipalpal identity; by comparison, knockdown of the spatially restricted paralog *Deformed-A* in the spider affects only the first walking leg. To investigate the genetic basis for leg elongation and tarsomere patterning, we identified and interrogated the function of an *Epidermal growth factor receptor (Egfr)* homolog. Knockdown of *Egfr* incurred shortened appendages and the loss of distal leg structures. The overlapping phenotypic spectra of *Egfr* knockdown experiments in the harvestman and multiple insect models are striking because tarsomeres have evolved independently in these groups. Our results suggest a conserved role for *Egfr* in patterning distal leg structures across arthropods, as well as cooption of EGFR signaling in tarsomere patterning in both insects and arachnids. The establishment of genomic resources for *P. opilio,* together with functional investigations of appendage fate specification and distal patterning mechanisms, are key steps in understanding how daddy-long-legs make their long legs.

## Introduction

Arthropods represent an ancient and asymmetrically species-rich phylum of animals. Within this group, the subphylum Chelicerata (e.g., spiders, scorpions, mites, horseshoe crabs, etc.) constitutes the sister group to the remaining Arthropoda, making it a key lineage for deciphering macroevolutionary phenomena common to the group, such as the evolution of segmentation and tagmosis, and the nature of the ancestral jointed appendage. Moreover, many chelicerate groups exhibit evolutionary novelties that are thought to play a role in the diversification of their constituent lineages, such as complex silks (spiders), venoms (spiders, scorpions, ticks, and pseudoscorpions), and acute vision (jumping spiders).

Developmental genetic and genomic resources for the comparative biology of chelicerates have largely prioritized spider, mite, and tick models. These resources have revealed complex dynamics in the evolution of chelicerate genomes. A group of six terrestrial orders (Arachnopulmonata), which includes spiders, scorpions, and pseudoscorpions, exhibit an ancient (pre-Devonian), shared whole genome duplication, as evidenced by the number and architecture of Hox clusters, intraspecific analysis of synteny, patterns of microRNA enrichment, gene expression patterns, and gene tree topologies (Leite et al., 2018; 2016; Nolan, Santibáñez López, & Sharma, 2020; Ontano et al., 2020; Schwager et al., 2017; Sharma, Schwager, Extavour, & Wheeler, 2014b) (Fig. 1*A*). Separately, genomes of all four living Xiphosura (horseshoe crabs) suggest a lineage-specific twofold genome duplication in this order, with one of these duplications occurring relatively recently (Battelle et al., 2016; Kenny et al., 2015; Roelofs et al., 2020; Shingate, Ravi, Prasad, Tay, & Venkatesh, 2020a; Shingate et al., 2020b). While genomes of Acariformes and Parasitiformes (mites and ticks) suggest that these two orders were not included in the genome duplication events, their genomes deviate from those typical of most arthropods. As examples, the acariform mite *Tetranychus urticae* exhibits extreme genome compaction (90 Mb), in tandem with loss of many transcription factors, which has been linked with miniaturization in this species (Grbić et al., 2011). Similarly, the genome of the parasitiform mite *Galendromus occidentalis* has shown atomization of Hox clusters, degradation of synteny, and comparatively high rates of intron gain and loss (Hoy et al., 2016). Such genomic trends parallel the accelerated evolution of these two orders across phylogenetic datasets.

**Fig. 1.**
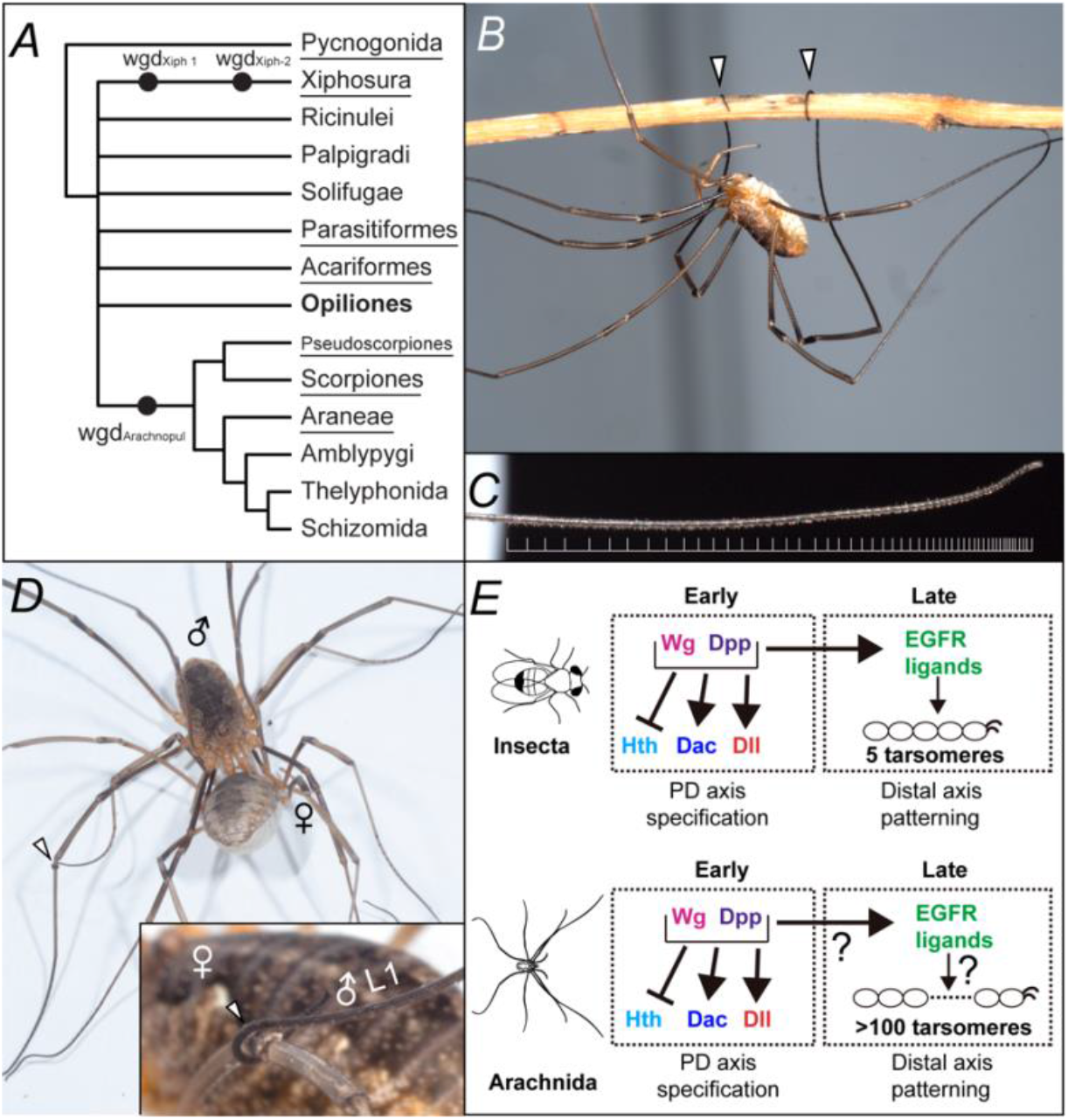
The significance of Opiliones in evolutionary developmental biology. (*A*) Consensus phylogeny of Chelicerata (based on (Ballesteros & Sharma, 2019; Lozano-Fernandez et al., 2019; Ontano et al., 2020; Sharma, Kaluziak, Pérez-Porro, González, Hormiga, et al., 2014a)), and inferred whole genome duplication (WGD) events in Xiphosura and Arachnopulmonata. (*B*) Adult male *P*. *opilio* climbing on a twig using its prehensile tarsi. (*C*) Detail of the distal subdivisions (tarsomeres) of the leg 2 tarsus. The distal terminus is to the right. White bars mark tarsomere boundaries. (*D*) Male *P. opilio* (above) copulating with a female (below). Note the use of prehensile tarsus to hold the female’s legs (arrowhead). Inset: Male tarsus holding female’s leg after mating. (*E*) Gene regulatory network (GRN) specifying PD axis and distal appendage patterning in insects and arachnids. While the Wg-Dpp GRN role in patterning PD axis is conserved in Arthropoda, no functional data exist informing EGFR signaling in non-insect arthropods. Pictures: Caitlin M. Baker.

One group that may facilitate comparative genomics of Chelicerata is the arachnid order Opiliones (harvestmen) (Fig. 1*B*). In phylogenomic datasets, Opiliones as a group exhibit evolutionary rates that are lower than those of Parasitiformes or Acariformes, and their placement outside of arachnopulmonates makes this group phylogenetically significant (Ballesteros & Sharma, 2019; Lozano-Fernandez et al., 2019; Sharma, Kaluziak, Pérez-Porro, González, Hormiga, et al., 2014a). Developmental transcriptomes of the emerging model species *Phalangium opilio* have suggested that harvestmen do not exhibit systemic genome duplication, as evidenced by Hox gene complements (Sharma, Schwager, Extavour, & Giribet, 2012), absence of paralogy across the homeobox gene family (Leite et al., 2018), and gene expression patterns of single-copy homologs of appendage-patterning genes that exhibit paralogs with divergent expression patterns in arachnopulmonates (Leite et al., 2018; Nolan et al., 2020). As a result, *P. opilio* has proven useful for study of chelicerate developmental biology, from the perspective of comparative gene expression as well as testing the conservation of gene regulatory networks across Arthropoda.

Beyond its use in polarizing developmental mechanisms on phylogenetic trees, Opiliones also exhibit a suite of unique characteristics that are not found in other panarthropod model systems. The most salient of these is the incidence of elongate walking legs in some groups (e.g., the superfamily Phalangioidea, commonly termed “daddy long legs”). Beyond the hypertrophied growth of certain leg segments, most harvestmen exhibit subdivision of the tarsus into pseudosegments called tarsomeres, with over 100 tarsomeres in some Phalangioidea (Shultz & Pinto-da-Rocha, 2007) (Fig. 1*C*). Tarsomeres have evolved repeatedly across the arthropod tree of life, as exemplified by derived insects (Kojima, 2017), scutigeromorph centipedes (Kenning, Müller, & Sombke, 2017), and several arachnid orders. However, the tarsomeres of harvestmen are sufficiently numerous in some species that they confer prehensility to the distal leg, which is known to be used in behaviors like climbing vegetation, male-male combat, and courtship (21-23) (Fig. *1B*, 1*D*). Moreover, the number of tarsomeres varies across the legs of many harvestmen, which has facilitated the derivation of a new appendage function; the largest number of tarsomeres in Phalangioidea typically occurs on the antenniform second leg pair, which is mostly used for sensing the surroundings and not for walking (Willemart, Farine, & Gnaspini, 2009).

While these aspects of Opiliones biology make them an opportune group for investigation of comparative genomics and gene function, neither a genome nor functional data on the developmental basis for leg elongation are available in this group. With respect to tarsomere patterning, functional genetic data do not exist for non-insect arthropod groups. Here, we generated a genome assembly for the model system *P. opilio,* with the aim of characterizing their genome architecture with respect to better studied chelicerate groups. To test the hypothesis that single-copy homologs of harvestmen exhibit a broader ancestral function that was subsequently subfunctionalized in the two paralogs of arachnopulmonates, we interrogated the function of the Hox gene *Deformed (Dfd)* using RNA interference (RNAi) in the harvestmen. As a first step in deciphering how daddy long legs make their eponymous appendages, we focused on the most distal segment of the leg, the tarsus. In holometabolous insects, tarsal fate is under the control of the Epidermal Growth Factor Receptor (EGFR) signaling pathway (G. Campbell, 2002; Galindo, Bishop, Greig, & Couso, 2002; Grossmann & Prpic, 2012; Kojima, 2004; Nakamura, Mito, Miyawaki, Ohuchi, & Noji, 2008) (Fig. 1*E*). We therefore identified an *Egfr* homolog in *P. opilio* and assessed its function in tarsal patterning.

## Results

### *General features of* P. opilio *draft genome assembly*

The draft assembly of the *P. opilio* genome (2n = 24) (Jindrová, Hirman, Sadílek, Bezdecka, & Stahlavsky, 2020) comprises 580.4 Mbp (37.5% GC content) in 5137 scaffolds (N50: 211089) and 8349 contigs (N50: 127429; Fig. S1; Table S1). The assembled genome size is slightly larger than the estimated genome size based on the analyses of the short read sequencing data in GenomeScope (465 Mbp) (Vurture et al., 2017) (Fig. S2). The predicted genome repetitiveness is 54.4% and has an estimated heterozygosity of 1.24%. The repetitiveness is further corroborated by the output of RepeatMasker, which indicated 46.85% of the genome consisted of repetitive sequences. The number of predicted genes is 20,315, which was further refined with a 98% similarity threshold and manual curation to a final gene set of 18,036. This is comparable to predicted gene sets for ticks, *Ixodes scapularis* (20,486) and mites, *Tetranychus urticae* (18,414) (Grbić et al., 2011; Gulia-Nuss et al., 2016). An assessment using the arthropod set of benchmarking universal single-copy orthologs (BUSCO (Waterhouse et al., 2017)) indicates 95.1% completeness, single copy complete 88.0%, single copy duplicated 7.1%, with 2.3% fragmented and 2.6% missing (Fig. S1; Table S1). Contamination assessment based on sequence coverage and GC content supports a relatively contamination-free assembly (Fig. S3; Supplementary Methods).

### *The genome of* P. opilio *reveals the absence of Arachnopulmonata-specific whole genome duplications*

To assess whether *P. opilio* shares any of the whole genome duplication events affecting certain chelicerate lineages, we first examined the architecture of Hox clusters in this species. We discovered one 480 kb scaffold that bore six Hox genes, with the remaining four Hox genes occurring on smaller, individual scaffolds (Fig. 2*A* and *B*). In addition to the small size, these scaffolds contained very few or no adjacent genes, suggesting that the non-collinearity of the anterior Hox genes results from the fragmentation of the assembly. On the larger 480 kb scaffold, microRNAs *mir-10* and *iab-4* were located adjacent to *Dfd* and *abdA,* respectively, which reflect conserved positions with respect to other arthropods (Pace, Grbić, & Nagy, 2016) (Fig. 2*A*). The complete peptide sequences of all ten Hox genes discovered in the genome assembly corresponded to previous partial sequences predicted from developmental transcriptomes of this species (Sharma et al., 2012).

**Fig. 2.**
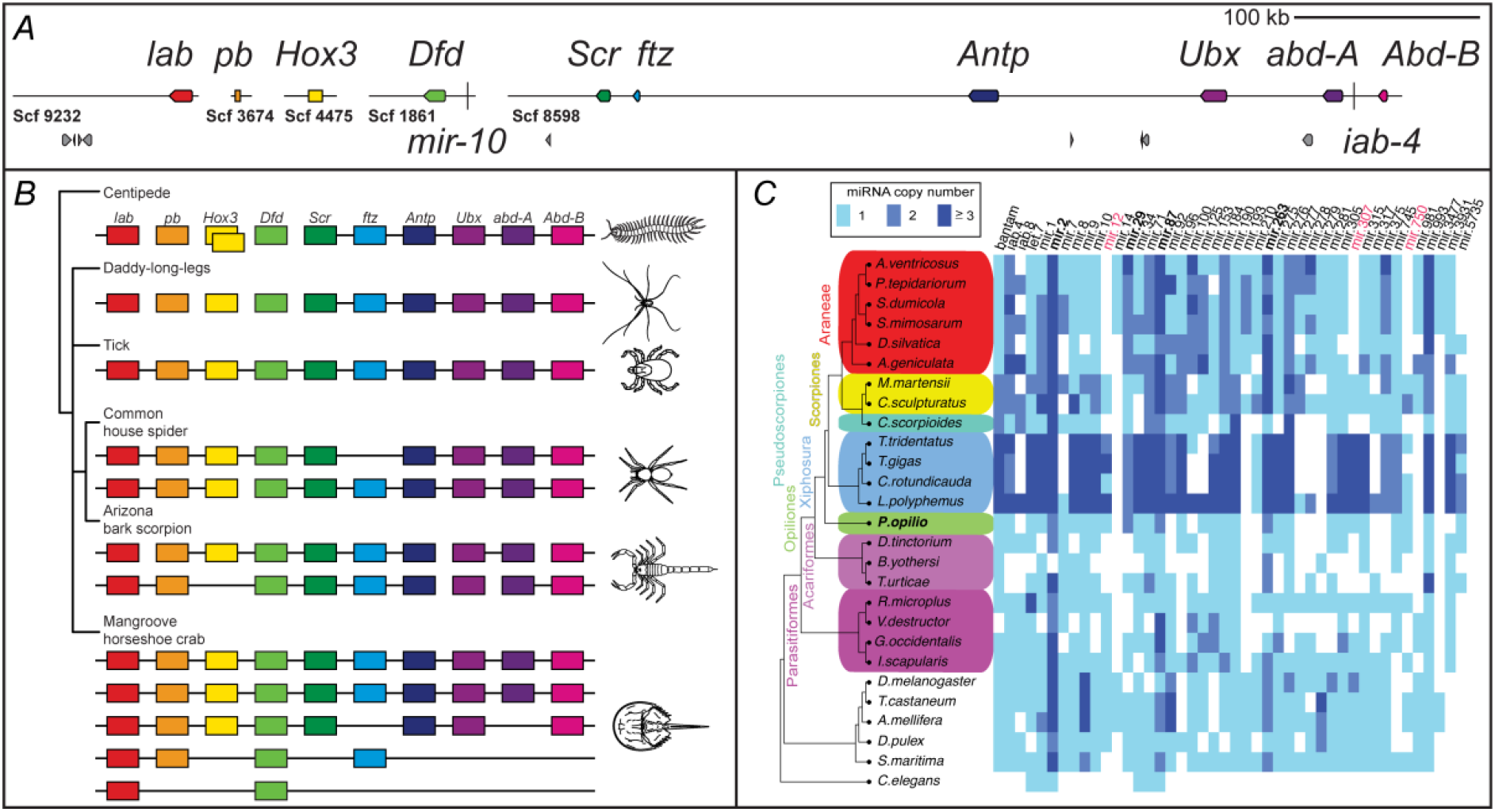
Hox genes and microRNAs support an unduplicated genome in the daddy-long-legs *P. opilio*. (*A*) Hox gene-containing scaffolds to scale. miRNAs *mir-10* and *iab-4* are represented by vertical bars. Hox genes are depicted in colored boxes, and other predicted genes in grey boxes. (*B*) Hox clusters in selected arthropod genomes (after (Chipman et al., 2014; Gulia-Nuss et al., 2016; Schwager et al., 2017; Shingate et al., 2020b)). (*C*) Comparative analysis of miRNA families and ortholog copy numbers in *P. opilio* and other chelicerates supports retention of single copies of several families in harvestmen, in contrast with duplication found in arachnopulmonates. Columns correspond to individual miRNA families, with colors representing different number of paralogs. miRNAs in bold are duplicated in *P. opilio* and most other chelicerates. Red text indicates families absent in spiders, but present in scorpions, pseudoscorpions and harvestmen.

Through manual annotation, we separately examined the genome for evidence of duplicates in genes with known arachnopulmonate-specific paralogs and distinguishable embryonic gene expression patterns, focusing on four leg patterning genes *(dachshund* (Battelle et al., 2016), *homothorax* (Gong et al., 2019), *extradenticle* (Schwager et al., 2017), *spineless* (Setton et al., 2017)) and the retinal determination gene network *(sine oculis, Optix, orthodenticle* (Gainett et al., 2020; Samadi, Schmid, & Eriksson, 2015; Schomburg et al., 2015)). All of these genes, known to exhibit arachnopulmonate-specific duplications and spatiotemporal subdivision of expression patterns, were discovered as single-copy in the harvestman genome (Table S2).

A prediction of a recent WGD event is that a large proportion of paralogous portions of the genome (paranomes) shares the same age. We assessed whether the *P. opilio* genome shows signal of recent genome expansion by comparing the Ks values (an estimate of synonymous distance between paralogs as a function of divergence time; used for detection of recent WGD events) with those of the genomes of selected arthropods. In accordance with a recent study (11), we corroborated a peak of Ks values in two horseshoe crab genomes, which suggests that at least one of the WGD duplications in this order is relatively recent. Ks plots did not show peaks in the arachnopulmonates, in accordance with the ancient timing inferred for this event (>500 My) (Fig. S4). We also found no evidence of a recent systemic duplication in *P. opilio* (Fig. S4).

We next examined the distribution of families of microRNAs (miRNAs), which are noncoding RNAs about 22 bp long, with important regulatory roles in animals. miRNAs have been used as effective phylogenetic characters (L. I. Campbell et al., 2011; Rota-Stabelli et al., 2011; Tarver et al., 2013; 2018; B. M. Wheeler et al., 2009) (but see (Thomson, Plachetzki, Mahler, & Moore, 2014)), and have been shown to exhibit the signature of genome duplication in both Arachnopulmonata and Xiphosura (Leite et al., 2016; Ontano et al., 2020). We searched for 44 selected miRNAs in the genome of *P. opilio* and seven chelicerate genomes not previously sampled for miRNAs, compiling a final dataset with 21 chelicerates and six outgroups (Table S3). Thirty conserved miRNA families were identified in the *P. opilio* genome, with 33-41 found in the six spiders, 36-40 in the two scorpions, 29 in the pseudoscorpion, 31-39 in the four xiphosurans, 17-27 in the mites, and 22-37 in the ticks (Fig. 2*C*). Among them, only families mir-2, mir-29, mir-87, and mir-263 had two homologs in Opiliones (Fig. 2*C*). Nonetheless, these four microRNAs are also duplicated in most other chelicerates, with the exception of mir-29 (only one homolog in Panscorpiones, Acariformes and Parasitiformes) and mir-263 (only one homolog in Pseudoscorpiones) (Fig. 2*C*). We found no miRNAs unique to harvestmen, arachnids, or arachnopulmonates. Lastly, families mir-12, mir-307, and mir-750 were found in most chelicerate orders, but not in spiders. Consistent with the inference of an unduplicated harvestman genome, we found no evidence of miRNA duplications in the harvestman genome that were shared either with Arachnopulmonata or Xiphosura.

### *The single-copy homolog of the Hox gene* Deformed *patterns the identity of two leg-bearing segments in the harvestman*

*Deformed* homologs of arachnids are expressed in the leg-bearing segments (L1–L4), and, in the case of mites, expression also extends posteriorly (Schwager et al., 2017; Sharma et al., 2012; Telford & Thomas, 1998). In *P. tepidariorum,* an arachnopulmonate inferred to have undergone a WGD, the two *Dfd* paralogs have divergent expression patterns, with only *Ptep-DfdA* being expressed in the legs (Schwager et al., 2017) (Fig. 3*A*). *Ptep-DfdA* knockdown via RNA interference (RNAi) results in a L1-to-pedipalp homeotic transformation (Pechmann, Schwager, Turetzek, & Prpic, 2015) (Fig. 3*B* and *C*). No functional data exist for the single-copy *Dfd* homolog of non-arachnopulmonate arachnids (the plesiomorphic condition). We investigated the function of the single-copy *Dfd* homolog of *P. opilio (Popi-Dfd*) through RNAi, via embryonic injection of double-stranded RNA (dsRNA) (Fig. S5). Upon completion of embryogenesis, 56.6% *(Popi-Dfd5’* fragment; n=43/76) to 24.7% *(Popi-Dfd3* fragment; n=43/174) of dsRNA-injected embryos exhibited leg-to-pedipalp homeotic transformation, a phenotype never observed in negative control embryos (Fig. S5). Strongly affected embryos from *Popi-Dfd* dsRNA treatment were able to hatch and underwent complete L1-to-pedipalp homeotic transformation (Fig. 3*D* and *E*), as evidenced by the loss of the metatarsus (an arachnid leg-specific segment, absent in pedipalps), the presence of pedipalp-specific setal spurs, and the loss of the leg-specific tooth of the distal claw (Fig. *3F–G* and *I–J*). In contrast to the spiders exhibiting *DfdA* loss-of-function phenotypes, strongly affected daddy-long-leg RNAi embryos (class I) also exhibited a L2-to-pedipalp transformation, as evidenced by the absence of the metatarsus; the presence of distal pedipalp setal pattern, and the loss of the distal claw tooth (Fig. 3*H* and *K*). transformed L2 also exhibited a fusion of tibia and patella segments (Fig. 3*K*). Weakly affected embryos (class II) underwent only partial L1-to-pedipalp transformation (absence of metatarsus) (Fig. 3*M*), and exhibited normal L2 phenotype, sometimes with a notched L2 metatarsus. The majority of *Popi-Dfd* knockdown phenotypes exhibited mosaicism, with one side of the body more strongly affected (bilateral mosaics), likely due to localization of dsRNA at the site of injection (Fig. 3*L*). Reduced *Popi-Dfd* expression was observed overall in embryos injected with *Popi-Dfd* dsRNA (Fig. S6) and reduced expression correlated strongly with the side of the body presenting homeotic transformation in mosaic embryos (Fig. S6).

**Fig. 3.**
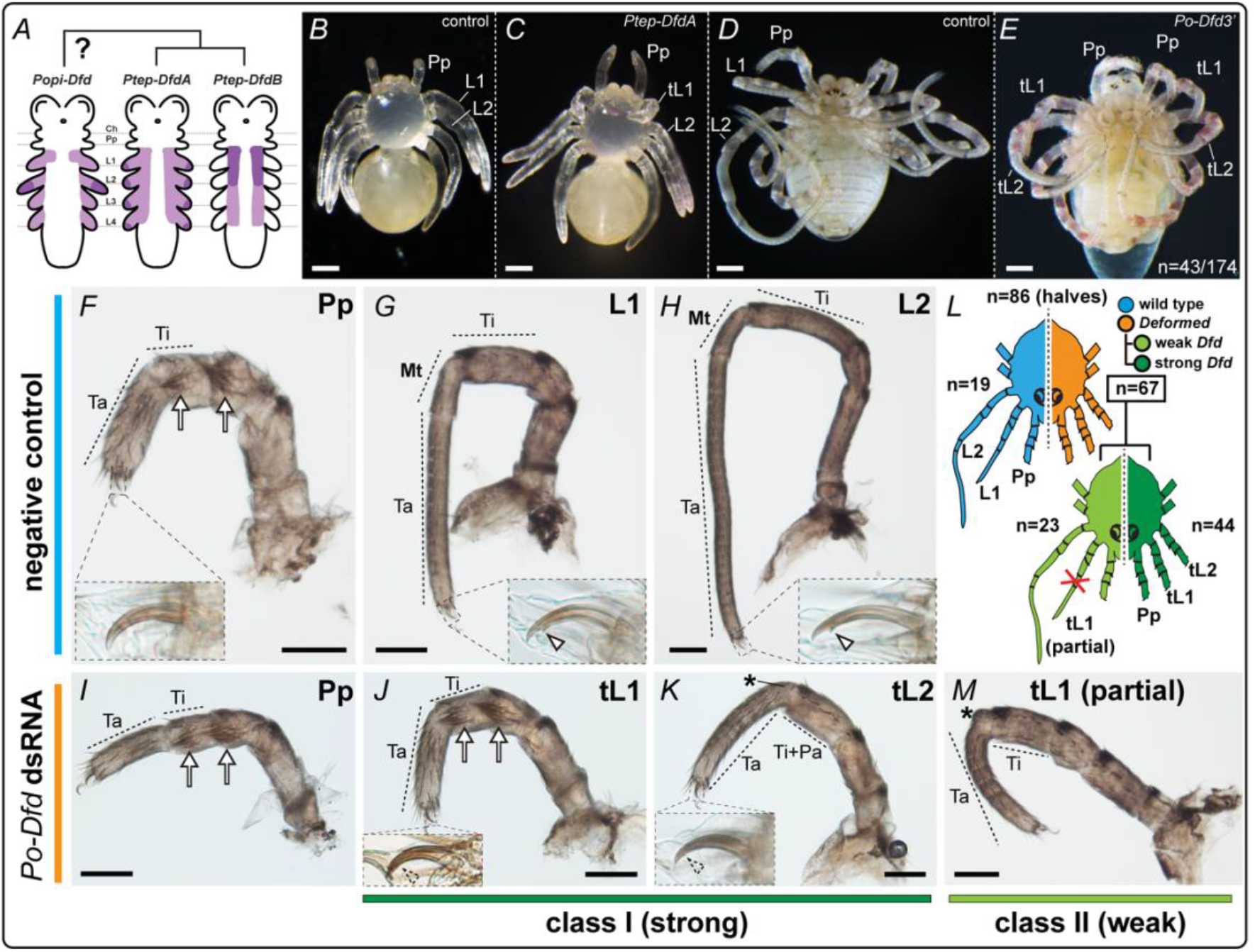
The single-copy ortholog of *Deformed* in *P. opilio* represses anterior appendage identity in two body segments. (*A*) Schematic of expression patterns of *Deformed* homologs in the germ band of the daddy-long-legs (*P. opilio*) and the common house spider (*P. tepidariorum*). Light: weak expression; dark: strong expression (*B*) Control hatchling (wild type) of *P. tepidariorum,* ventral view. (*C*) *P. tepidariorum* hatchling from the *Pt-DfdA* dsRNA-injected treatment. The first pair of legs is homeotically transformed into pedipalps, whereas posterior legs (L2-L4) are not affected. (*D*) Wild type (negative control) hatchling of *P. opilio.* (*E*) *P. opilio* hatchling from *Po-Dfd* RNAi treatment. The first and second pairs of legs are homeotically transformed into pedipalps. (*F-H*) Appendage mounts of wild type *P. opilio* hatchlings in lateral view (in order: pedipalp, L1 and L2, and tarsal claws (insets)). *(IK* and *M)* Appendage mounts of *Po-Dfd* RNAi hatchlings in lateral view. *(I)* Pedipalp. *(J)* Class I phenotype (strong) showed L1-to-pedipalp transformation. Note pedipalp-specific spurs, absence of claw tooth (inset), and absence of metatarsus (penultimate segment). (*K*) Class I phenotype (strong) showed L2-to-pedipalp homeosis. Note the absence of claw tooth (inset) and reduced metatarsus (asterisk). The tibia and patella are fused in transformed L2. (*L*) Schematic depiction of the mosaicism observed in *Po-Dfd* RNAi hatchlings and distribution of phenotypic classes. (*M*) Class II phenotype (weak), partial leg 1 to pedipalp transformation. Asterisk, reduced metatarsus; arrow, setal spurs; arrowhead, claw tooth; L1, leg 1; tL1, transformed leg 1; L2, leg 2; tL2, transformed leg 2; Mt, metatarsus; Pa, patella; Pp, pedipalp; Ta, tarsus; Ti; tibia. Scale bars: 100 μm.

In summary, in contrast to the patterning of a single leg-bearing segment by the *DfdA* paralog in the spider, the single-copy *Dfd* homolog of *P. opilio* is necessary for conferring leg identity of two segments immediately posterior to the tritocerebral somite (the third head segment).

Epidermal growth factor receptor *is necessary for distal leg patterning in the daddy-long-legs* The proximal-distal (PD) axis of the appendages of Arthropoda is specified by a conserved gene regulatory network that involves the interactions of *wingless* (*wg*), *decapentaplegic (dpp)* and a suite of leg gap genes (e.g., *Dll, hth, dac)* (Angelini & Kaufman, 2005; Estella, Voutev, & Mann, 2012; Kojima, 2004; Pechmann, Khadjeh, Sprenger, & Prpic, 2010; Setton & Sharma, 2018). On the other hand, the fate of the most distal segment of the leg, the tarsus, is established later in development by a Wg-Dpp independent mechanism, at least in insects (G. Campbell, 2002; Galindo et al., 2002; Grossmann & Prpic, 2012; Kojima, 2004; Nakamura et al., 2008). In the fruit fly *Drosophila melanogaster*, an initial input of Wg and Dpp establishes a source of Epidermal Growth Factor Receptor (EGFR) signal in the distal leg territory, which then independently governs the formation of tarsal subdivisions (tarsomeres) by regulating several downstream tarsal genes (Fig 1 *E*) (G. Campbell, 2002; Estella et al., 2012; Galindo et al., 2002; Galindo, Bishop, & Couso, 2005).

It is not known whether the mechanism of distal leg patterning established for insects applies to other arthropods, such as chelicerates. We therefore leveraged genomic resources for *P. opilio* and other chelicerates to address the patterning of the tarsomeres, focusing on *Egfr.* We discovered two putative *Egfr* homologs in the *P. opilio* genome. Two copies of *Egfr* also occur in the pseudoscorpion *Conichochernes crassus*, and the arachnopulmonates *P. tepidariorum* (spider) and *Phrynus marginemaculatus* (whip spider) (Fig. S7). Multiple copies occur in the genome of horseshoe crabs (Fig. S7). Phylogenetic analysis showed that each paralog of the spider forms a clade with a paralog of the whip spider and the pseudoscorpion, suggesting that the paralogs derive from the shared duplication event at the base of Arachnopulmonata. All horseshoe crab *Egfr* copies formed a clade, suggesting a lineage specific duplication in this case (Fig. S7). The harvestman *Egfr* paralogs also formed a clade to the exclusion of other arachnids. These results accord with the independent wholegenome duplication in Xiphosura and Arachnopulmonata (with the inclusion of Pseudoscorpiones (Ontano et al., 2020)). and suggest that the harvestmen paralogs are the result of a lineage-specific duplication event.

*Popi-EgfrA* protein prediction contains the typical *Egfr* domains known from *D. melanogaster*, including the extracellular L-domains, Furin-like, and GF receptor IV, the transmembrane region, and the intracellular Tyrosine-Kinase domain (Fig. S8). On the other hand, *Popi-EgfrB* contains only the extracellular domains, and lacks the transmembrane and intracellular domains seen in the other metazoan *Egfr* (Fig. S8) (Barberán, Martín-Durán, & Cebrià, 2016). A 3’ UTR for *Po-EgfrB* was assembled in both embryonic transcriptomes and corroborated by the genome assembly, disfavoring fragmentary assembly as a possible explanation for missing domains.

Given the divergent nature of *Popi-EgfrB* and apparent absence of a tyrosine-kinase intracellular domain, we focused our functional investigation on the paralog containing all known *Egfr* functional domains *(Popi-EgfrA).* We first assayed expression of this paralog during harvestman embryogenesis using in situ hybridization. *Popi-EgfrA* is ubiquitously expressed in low levels in early germ band stage embryos (Stage 10). At this stage, *Po-EgfrA* is also expressed in a strong circular domain around the region of the future mouth (stomodeum), in a strong ring at the base of each appendage, and in a weak but well-defined broad stripe along the ventral midline (Fig. 4*A* and *A’*; *4D-G*). In later stages, additional expression domains occur in the developing eye field, in a strong domain in the medial bridge of the developing brain, and in rings at the boundaries of the segments (podomeres) of the developing appendages (Fig. 4*B* and *C*; 4*H-K*).

**Fig. 4.**
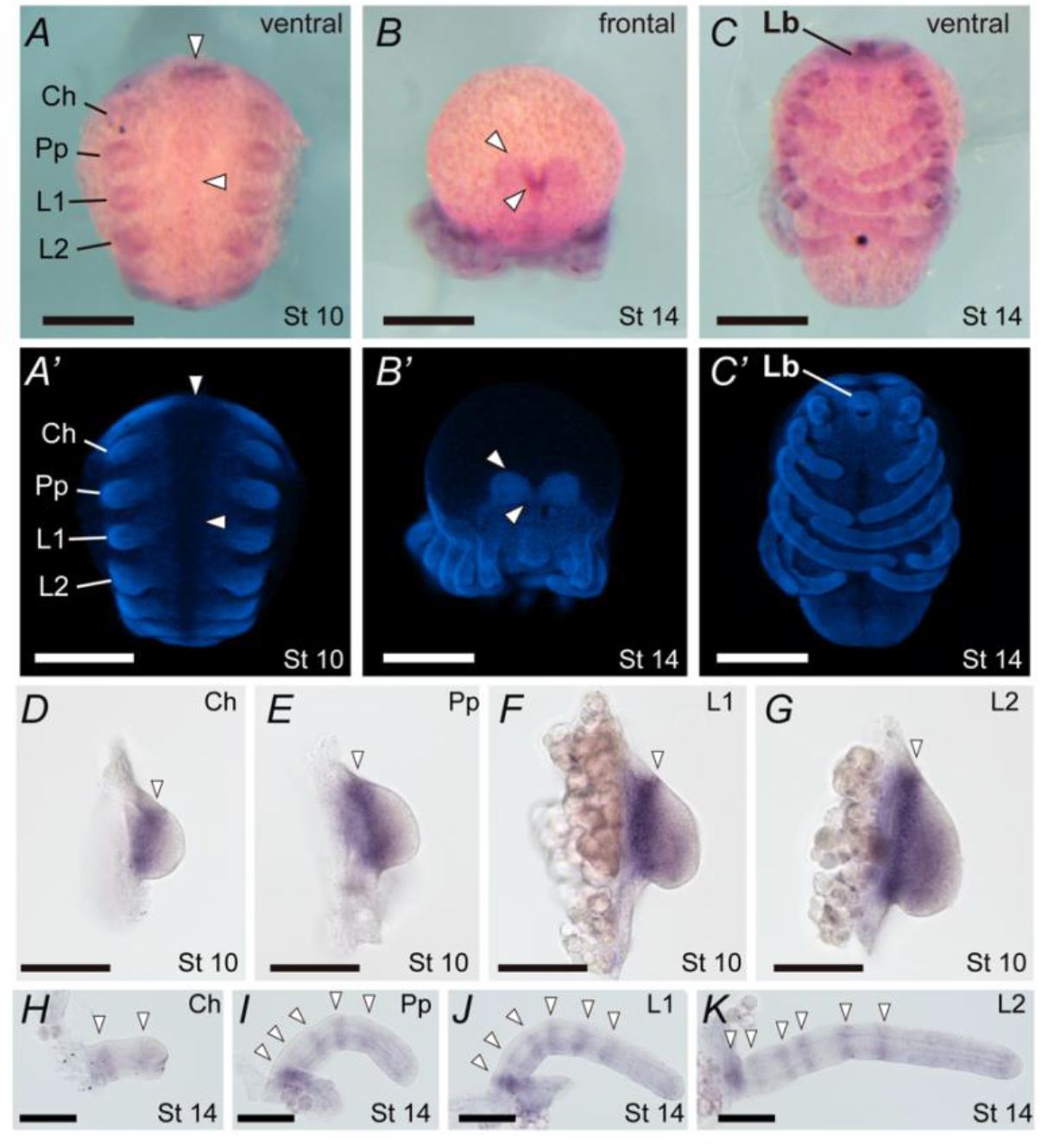
*Popi-EgfrA* in situ hybridization in wild type embryos shows conservation of expression domains in the daddy-long-legs with respect to insects. *A-C* and *D-K* in brightfield and *A’-C’* in Hoechst counterstain. (*A* and *A*’) Stage 10, ventral view, with domains of expression in the ventral midline, appendages, and stomodeum. (*B* and *B’)* Stage 14 frontal view, with expression on the head lobes and strong domain on the medial bridge of developing brain. (*C* and *C’*) Stage 14, ventral view. *(D-G)* Stage 10 appendage mount in lateral view of, respectively, chelicera, pedipalp, L1, and L2. *(H-K)* Stage 14 appendage mount in lateral view of, respectively, chelicera, pedipalp, L1, and L2. Arrowheads mark discreet domains of expression. Ch, chelicera; Pp, pedipalp; L1, leg 1; L2, leg 2. Scale bars: 200 μm (*A-C*); 100 μm (*D-K*).

To investigate the function of *EgfrA,* we performed embryonic RNAi against this paralog. RNAi against *EgfrA* resulted in 39.6% (n=36/91) of hatchlings exhibiting segmentation and appendage defects not observed in control embryos (Fig. 5) (Fig. S9); all affected embryos were bilateral mosaics (Fig. 5*B* and *C*), and defects correlated with reduced *Po-EgfrA* expression in embryos (Fig. S10). Hatchlings from *Popi-EgfrA* dsRNA treatment showed defects of antero-posterior (body) segmentation, with dorsal tissue fusion on the side of the body affected (n=29/36), correlating with a characteristic curved shape of the body of mosaic individuals (Fig. 5*B*). This segmentation phenotype accords closely with a *Egfr* loss-of-function phenotype in the beetle *T. castaneum* (Grossmann & Prpic, 2012). Defects of the eyes (n=25/36) ranged from a small reduction in size to complete absence (Fig. 5*C*), according with a previously established role for EGFR signaling in eye development of *D. melanogaster* (Halfar, Rommel, Stocker, & Hafen, 2001).

**Fig. 5.**
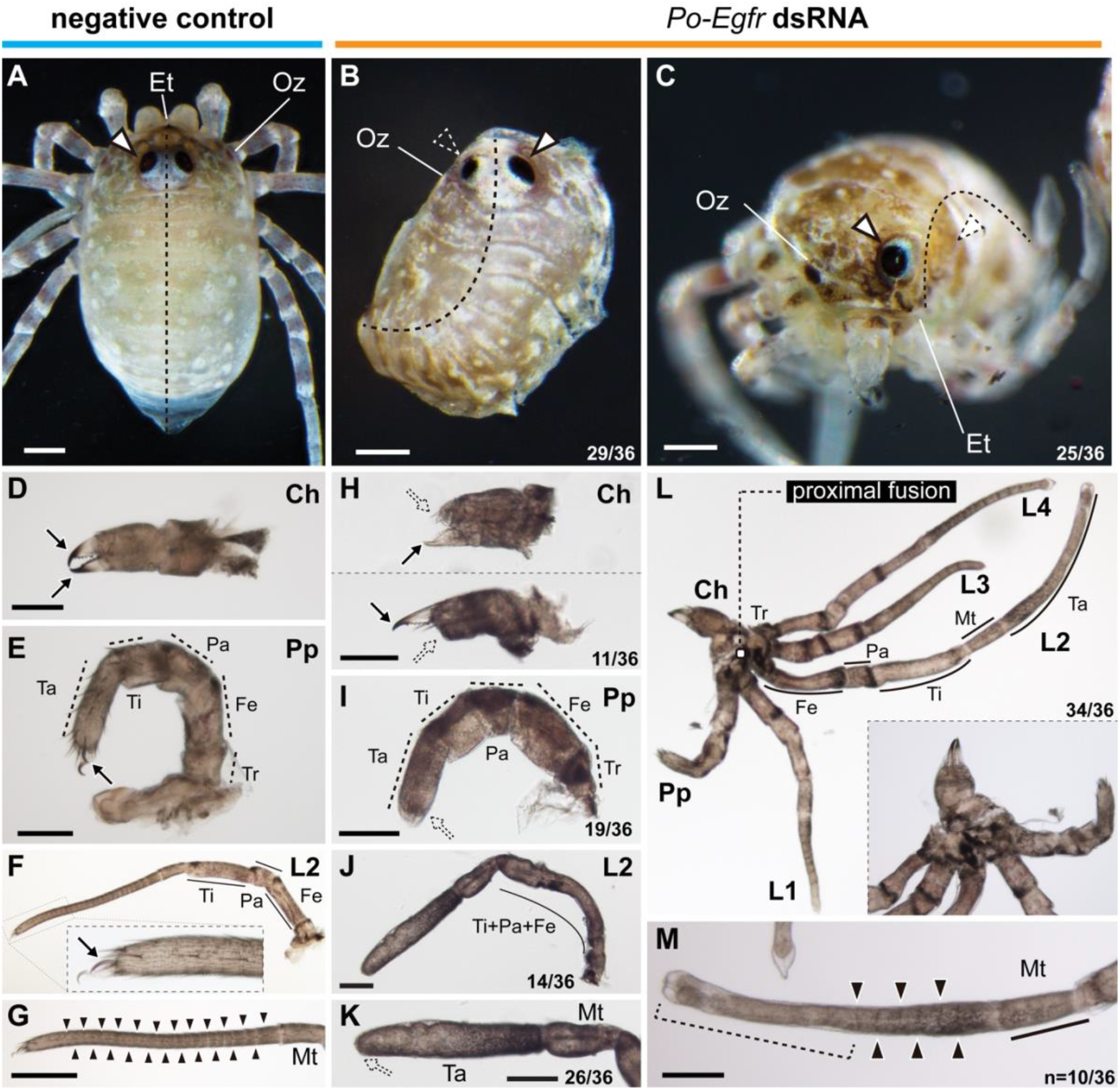
*Po-EgfrA* knockdown affects dorsal patterning, eyes, and appendage formation. (*A*) Negative control hatchling in dorsal view. (*B* and *C*) Hatchlings from *Po-EgfrA* dsRNA injected treatment (mosaic, left side affected). (*B*) Hatchling in dorsal view. Note dorsal fusion on the left side of the body (n=29/36). (*C*) Hatchling in frontal view, with the left eye absent. A subset of *Egfr* phenotypes showed eye reduction (25/36) *(D-G)* Appendage flat mounts of negative control hatchlings, in lateral view. (*D*) Chelicera. (*E*) Pedipalp. *(F)* L2. Inset shows detail of the claw at the distal leg. (*G*) Tarsus of L2. (*H-M*) Appendage flat mounts of hatchlings of *Po-EgfrA* dsRNA-injected treatment, in lateral view. (*H*) Chelicerae showed reduction of fixed finger (upper panel), movable finger (lower panel) or both (n=11/36). *(I)* Pedipalps lacked claws (n=19/36). (J) L2, exhibiting podomere fusions proximal to the tarsus (n=14/36). (*K*) Distal end of L2, exhibiting claw and tarsomere reduction (n=26/36). (*L*) Proximal fusion in adjacent appendages (Ch–L4) (n=34/36). Inset: Detail of fused coxae. *(M)* Tarsus of leg 2 shown in (*L*). Weakly affected legs lacked claws and distal tarsal joints (brackets) but retained proximal joints (n=10/36). Arrow, claw; outlined white arrowhead, eye; dotted white arrowhead, eye defect; solid black arrowhead, tarsomere joints; Ch, chelicera; Et, egg tooth; Fe, femur; L1 –L4, legs 1 –4; Mt, metatarsus; Oz, ozophore; Pa, patella; Pp, pedipalp; Ta, tarsus; Ti, tibia; Tr, trochanter. Scale bars: 100 μm.

The appendages of hatchlings from *Po-EgfrA* dsRNA-injected eggs had defects in terminal structures (Fig. 5*D*–*K*). The chelicera showed reduction of the fixed finger, movable finger, or both (n=11/36) (Fig. 5*H*); the pedipalp showed reduced claw (n=19/36) (Fig. 5*I*); and, in the case of all legs, the claw and tarsomeres were reduced (n=26/36) (Fig. 5*J* and *K*). Most notably, a subset of weakly affected individuals (n=10) showed a condition in which the distalmost tarsomeres were fused and the claw was missing, or just the claw was missing (Fig. 5*M*).

Segmental fusions in limbs occurred in a subset of hatchlings (n=14/36) (Fig. 5*J*). The *Po-EgfrA* expression at the limb segment boundaries and appendage segment fusions upon knockdown suggest a role in appendage segmentation. The proximal segment of the appendages (coxa) showed defects ranging from reduction in size to complete proximal fusion of adjacent appendage coxae (n=34/36) (Fig. 5*L*), which correlates with the strong ring of expression at the base of all appendages. The resulting coxal fusion and dorsal patterning defects upon knockdown, suggests that EGFR signaling is necessary for proper anteroposterior segmentation of the body. In two negative control embryos, we observed appendage malformation with fusion of adjacent appendages, indicating that a small percentage of appendage defects observed in *Po-Egfr* RNAi experiments may be due to off-target effects of the injection procedure. Nonetheless, given high incidence in RNAi experiments (n=34/36), and correlated strong expression of *Po-EgfrA* in the coxa of all appendages, fused coxae appear to be an on-target phenotype of *Po-EgfrA* dsRNA injection. These results are consistent with expression and functional data available for *Egfr* homologs in insect models (Grossmann & Prpic, 2012; Nakamura et al., 2008; Refki & Khila, 2015).

In sum, our genomic survey and functional experiments in *P. opilio* suggest that EGFR signaling has a conserved role in appendage patterning in the common ancestor of insects and arachnids (by extrapolation, across Arthropoda). With respect to the unique morphology of daddy-long-leg appendages, the phenotypic spectrum obtained by RNAi suggests that *EgfrA* may underpin the morphogenesis of the large number of tarsomeres observed across Opiliones

## Discussion

### RNAi in the daddy-long-leg informs the evolution of Hox genes in arachnids

In chordates, understanding developmental gene function in outgroup taxa that diverged before WGD within Vertebrata has been key to understanding the role of new genes in generating organismal diversity through processes like subfunctionalization and neofunctionalization of paralogs. Paralleling this trend, understanding the fate of ancient paralogs retained in derived groups like spiders and scorpions requires investigation of arachnids that diverged before the duplication event, such as mites, ticks and daddy-longlegs. We assembled and investigated the genome of the experimentally tractable harvestman species *P. opilio*, which revealed no evidence of systemic genome duplication events previously reported for arachnopulmonates and horseshoe crabs. Together with the genome architecture of model systems like *Ixodes scapularis* and *Tetranychus urticae,* the harvestman genome supports the notion that an unduplicated genome is the ancestral condition for arachnids. This reconstruction is further supported by the advent of recent genomic resources for Pycnogonida (Ballesteros et al., 2020) (sea spiders), the sister group to the remaining Chelicerata.

As a first step in understanding the function of single-copy Hox genes in arachnids, we investigated the function of *Dfd,* whose expression domain is associated with appendage patterning across Chelicerata. Subdivisions of expression domain of Hox duplicates in Arachnopulmonata are systemic, with Hox paralogs exhibiting spatial and temporal divergence in spiders (Schwager et al., 2017) as well as scorpions (Sharma, Schwager, Extavour, & Wheeler, 2014b). Whereas the spider paralog *Ptep-DfdA* is necessary for repressing pedipalp identity on a single body segment, L1, (Pechmann et al., 2015), our results show that the single copy *Popi-Dfd* is required for specification of both L1 and L2 identity. A two-segment-identity requirement for the anterior Hox gene *Dfd* has also been demonstrated in the hemipteran *Oncopeltus fasciatus*, the coleopteran *T. castaneum*, and the crustacean *Parhyale hawaiensis,* where the single-copy *Dfd* is necessary for the normal development of the mandibular and maxillary segments (Brown et al., 2000; C. L. H. A. T. C. Kaufman, 2000; Martin et al., 2016) – the positional homologs of the arachnid L1 and L2 segments (Hughes & Kaufman, 2002). This conserved function of the single copy *Dfd* observed between the three mandibulate species and the chelicerate *P. opilio* suggests that the patterning of these two positionally homologous segments is an ancestral function for Arthropoda, and that the function of *DfdA* in spiders reflects an arachnopulmonate-specific subdivision of the ancestral function.

Nevertheless, we note that possible alternative explanations for the difference in phenotypes reported for the spider and the daddy-long-legs could include the timing of the knockdown (maternal RNAi versus embryonic RNAi, respectively). In addition, the restriction of the *DfdA* loss-of-function phenotype to just the L1 segment in the spider may reflect compensation and rescue by its paralog, *Ptep-DfdB*, in the L2 segment. Notably, no RNAi experiments against *Ptep-DfdB* appear in the literature, as is the case with most of the spider Hox genes; this gap is attributable to the absence of phenotypic outcomes upon single-gene knockdown of many spider Hox genes using maternal RNAi. Likewise, investigations of paralog compensation and rescue in the spider model are hindered by the low penetrance of double-knockdowns using maternal RNAi in this species (Khadjeh et al., 2012). Testing hypotheses of spider Hox gene function likely requires the establishment of genome-editing techniques for *P. tepidariorum*.

### A conserved role for EGFR signaling in patterning tarsal identity in Arthropoda

The expression of an *Egfr* homolog and associated loss-of-function experiments in *P. opilio* substantiate a conserved role for this signaling pathway in patterning distal leg identity in the common ancestor of insects and arachnids, and by extension, across Arthropoda. While we expected a conserved role for EGFR in patterning distal leg structures broadly, we found remarkable correspondence in the EGFR loss-of-function phenotypic spectrum with respect to the patterning of tarsomeres of both insects and *P. opilio.* This result is notable because the tarsomeres of harvestmen and insects have evolved independently; an unsegmented tarsus is clearly the plesiomorphic state for both Opiliones and Hexapoda (Shultz & Pinto-da-Rocha, 2007; Tajiri, Misaki, Yonemura, & Hayashi, 2011). These findings raise the possibility that the interaction between EGFR and Notch signaling in patterning the joints of *D. melanogaster* may also be conserved, a hypothesis which is partially supported by the expression and function of Notch signaling pathway members in the spider *Cupiennius salei* (Prpic & Damen, 2009).

In *Drosophila melanogaster*, EGFR-Ras signaling is responsible for patterning the legs in two phases. First, EGFR signaling begins with a distal expression (central in the leg disc) of the EGFR ligand *vein* (vn), and of *rhomboid (rho),* a transmembrane protein required for cleavage and release of EGFR ligands, in the leg disc (G. Campbell, 2002; Galindo et al., 2002). Reduction of EGFR signaling at this early stage results in truncation of claws and distal tarsomeres (G. Campbell, 2002; Galindo et al., 2002; 2005). Stronger reduction of EGFR signaling results in progressively greater deletion of more distal leg structures, which is consistent with a distal-to-proximal requirement gradient of EGFR signaling by downstream tarsal patterning genes, and with a distal source of EGFR signaling (G. Campbell, 2002; Galindo et al., 2002; Kojima, 2004). A distal source of EGFR signaling is also in accordance with the distal-to-proximal requirement for EGFR signaling in the regenerating leg of the cricket *G. bimaculatus* (Nakamura et al., 2008). In *P. opilio,* 72% (n=26/36) of the hatchlings resulting from RNAi against *Egfr* exhibited defects in the tarsus. These defects ranged from absence of claws and distal tarsomere fusions, to complete failure to form all tarsal subdivisions. The spectrum of increasingly severe defects from distal-to-proximal observed upon *Egfr* knockdown in *P. opilio* suggests that daddy-long-leg tarsomeres are patterned by a gradient of EGFR signaling similar to *D. melanogaster* and *G. bimaculatus*. We infer that the variance in penetrance in our experiment is related to the localization of dsRNA at the site of embryonic injection and small differences in the volume injected per egg, as further evidenced by the large number of mosaic embryos observed (with severity of loss-of-function phenotype correlated with decreased expression (Fig. S10).

In a second phase of EGFR leg patterning in *D. melanogaster, vn* and *rho* are expressed as rings at the boundaries of embryonic tarsomeres (Galindo et al., 2005). Reduction or upregulation of EGFR signaling at this later stage in *D. melanogaster* results in defects on median tarsomeres and in failure to correctly pattern the tarsal joints (G. Campbell, 2002; Galindo et al., 2002). This second function occurs via repression of components downstream of *Serrate* in the Notch signaling pathway (Galindo et al., 2005), which are responsible for the position and morphogenesis of joints (Estella et al., 2012; Kojima, 2004). Aside from tarsomere defects, triple null mutants for EGFR components *vn*, *rho* and *Roughoid* (which phenocopy *Egfr* down-regulation) also showed tissue defects proximal to the tarsus (G. Campbell, 2002). It was later proposed that effects proximal to tarsomere 3 are due to secondary signaling events or cell death (Galindo et al., 2005). However, in the short germband insects *T. castaneum*, *L. dissortis* and *G. bimaculatus* (Grossmann & Prpic, 2012; Nakamura et al., 2008; Refki & Khila, 2015), *Egfr* is expressed as rings at the boundaries of all leg segments proximal to the tarsus, in contrast to rings restricted to the tarsus in the fruit fly. As predicted from this expression pattern, RNAi-mediated knockdown in the first two species resulted in leg segment truncations proximal to the tarsus, of which at least in *T. castaneum* a TUNEL cell death assay disfavored an indirect role of cell death as driver of the leg-truncation phenotype (Grossmann & Prpic, 2012).

Our data in the arachnid *P. opilio* largely accord with the expression and functional results in short germ-band insects: 38% of *Po-EgfrA* RNAi phenotypes also exhibited leg segment fusions proximal to the tarsus. A notable difference is the stronger ring of expression on the coxa of all appendages, and resulting fusion upon RNAi, a phenotype not previously reported in the insect systems. Altogether, these results suggest that the second role of EGFR signaling in leg segmentation is also conserved in *P. opilio.* The smaller frequency of appendage segmental fusions observed in *P. opilio*, and concomitant strong tarsal defects in all these cases, may be the result of extended transcript degradation in cases of high penetrance. Disentangling the effects of early versus late EGFR signaling phases in the phenotypes observed in *Po-EgfrA* RNAi could be further explored by affecting EGFR signaling only later in development, to surpass early defects in PD axis development. This could be achieved by injecting dsRNA into embryos at later stages of leg development, with the prediction that leg segment joint defects would become more frequent than distal tarsomere defects (patterned by the first phase); or through the development of advanced functional approaches and transgenic tools. We anticipate that the genome of *P. opilio* will facilitate future investigations of developmental genomics in this model system.

## Materials and Methods

### Animal Husbandry

Founder population specimens of *P. opilio* were collected in Madison, Wisconsin, USA (43.074628, −89.403904), and a colony was maintained as previously described (Sharma et al., 2012). Full-sibling crosses were performed for six generations. Four fourth-generation adult male specimens and one sixth-generation female were immediately preserved in 95% ethanol. Prior to RNA and DNA extraction, samples were kept at −80°C. All specimens were deposited in the Smithsonian Institution National Museum of Natural History (USNM-ENTO145186-90). Three fourth generation male specimens are deposited as genomic vouchers at the Smithsonian Institution National Museum of Natural History (USNM-ENTO145186-188).

### RNA Sequencing

Total RNA was extracted from ca. 250 uL of *Phalangium opilio* embryos spanning an array of stages, reared from females captured in Weston, Massachusetts. RNA extraction was performed using TRIzol reagent (ThermoFisher), following the manufacturer’s protocol. mRNA was purified with the Dynabeads mRNA Purification Kit (Invitrogen), following the manufacturer’s protocol. cDNA libraries were constructed in the Apollo 324 automated system using the PrepX mRNA kit (IntegenX) and paired-end sequencing was performed at the Center for Systems Biology (Harvard University) on an Illumina HiSeq 2500 platform with a read length of 150 base pairs.

The resulting 79,472,462 paired end reads (NCBI PRJNA690950) were combined with an older library (16,225,145 paired end reads sequenced on an Illumina GA II; NBCI SRR1145735) for annotation of protein coding regions.

### Genome Sequencing

Total genomic DNA was isolated from two specimens (fourth-generation male USNM-ENTO145189 & sixth generation female USNM-ENTO145190). High molecular weight (HMW) genomic DNA was extracted using the Qiagen MagAttract HMW DNA Kit from 23-25 mg of snap-frozen tissue according to the manufacturer’s protocol. HMW Genomic DNA was quantified by fluorometry (Qubit, Thermo Fisher Scientific, Waltham, U.S.A.) and assessed for size by 1.0% (w/v) agarose gel using Pulse Field Electrophoresis (PFGE) agarose gel electrophoresis (Chef Mapper XA, Bio-Rad Laboratories, Inc).

For long-read sequencing, Single Molecule Real Time (SMRT) bell libraries were prepared according to the 20 kb Template Preparation Using BluePippin Size-Selection System (Pacific Biosciences) from HMW DNA of the fourth generation male (USNM-ENTO145189). After damage-repair the libraries were size-selected on a BluePippin system (0.75% (w/v) agarose gel cassette, dye-free, S1 marker, high pass 20kb protocol) to remove library fragments smaller than 10kb. Then libraries were recovered by PB AMPure beads, quantified by the high sensitivity fluorometric assay (Qubit, Thermo Fisher Scientific, Waltham, U.S.A.) and quality assessed using the genomic assay on the TapeStation (Agilent, Waldbronn, Germany). The prepared library was run on the PacBio Sequel platform (Pacific Biosciences) using version 2.1 chemistry. The Single Molecule Real-Time (SMRT) Cells were sequenced on (16) SMRT cells with 360min movie lengths. Long reads are deposited in NCBI (SRR12286133).

For short read sequencing, paired-end sequences were generated using HMW DNA from the sixth-generation female (USNM-ENTO-145189) on one lane of an Illumina HiSeq 2500 with a 350-bp insert size. Library preparation and sequencing of short reads were conducted at Novogene Co., Ltd. Short read sequences are deposited on NCBI (accession number SRR12286133).

### Genome Assembly and Annotation

Detailed methods and commands for genome assembly and annotation are provided in the Supplementary Material. Briefly, long reads were assembled using Canu v. 1.7 (Koren et al., 2017) with default parameters. The resulting contigs from Canu were processed using two rounds of scaffolding with SSPACE-LongRead v1.1 (Boetzer & Pirovano, 2014), followed by gap filling with PBJelly v. 15.8 (English et al., 2012) and further polishing with Pilon v. 1.23 (Walker et al., 2014). Haplotypic duplicates were identified and removed with Purge_dups v1.2.3 (Guan et al., 2020). Prior to annotation, a custom repeat library was constructed using RepeatModeler open-1.0.11 (Smit, Hubley, & Green, n.d.). Identified repeats were masked with RepeatMasker open-4.0.6. (Smit et al., n.d.). Gene predictions using BRAKER2 v.2.1.5 (Hoff, Lomsadze, Borodovsky, & Stanke, 2019) used previously generated RNA-Seq reads from developmental transcriptomes (NCBI SRX450969; (Ballesteros & Sharma, 2019; Sharma, Kaluziak, Pérez-Porro, González, Hormiga, et al., 2014a)). RNAseq reads were mapped to the masked genome using HISAT2 v. 2.2.0 (Kim, Paggi, Park, Bennett, & Salzberg, 2019) using default parameters. Gene predictions were validated using BLASTx and Blast2GO (Götz et al., 2008). Orthologs of interest (e.g., Hox and appendage patterning genes) were refined from BRAKER2 predictions by manually inspecting alignments containing scaffolds and the transcripts of interest retrieved from two embryonic transcriptomes of *P. opilio* previously published by us (NCBI SRX450969). Genome profiling (size estimation, heterozygosity, repetitiveness) was conducted in GenomeScope 2.0 (Vurture et al., 2017) using short read sequence data. The final assembly was subjected to contamination screening in Blobtools v.1.0 (Laetsch & Blaxter, 2017). We used Benchmarking Universal Single-Copy Orthologs (BUSCO v4.0.2 (Waterhouse et al., 2017)) to assess the assembly completeness by comparing it to 1,013 orthologs contained in the arthropoda_odb10 gene database (Kriventseva et al., 2019). The genome assembly was deposited on NCBI (PRJNA647749) and a genome browser was generated using MakeHub (Hoff, 2019).

### Orthology Inference and Phylogenetic Analysis

Orthologs in the embryonic transcriptomes and genomes of species used in phylogenetic analyses were identified with tBLASTn (Altschul, Gish, Miller, Myers, & Lipman, 1990). All unique blast hits were retained, translated to protein using TransDecoder v.5.5.0 (Haas et al., 2013), and clustered with a similarity threshold of 95% using CDHIT 4.8.1 (Fu, Niu, Zhu, Wu, & Li, 2012). Preliminary alignments were inferred with Clustal Omega (Sievers et al., 2011). Preliminary phylogenetic analyses were conducted with approximate maximum likelihood in FastTree v. 2.1.10 (Price, Dehal, & Arkin, 2010) to selected candidates forming a clade with BUSCO arthropod orthologs of interest.

For *Egfr* searches we used as queries protein sequences from *Drosophila melanogaster* (FBpp0071570), *Tribolium castaneum* (XM_008193271.2), *Limnoporus dissortis* (KF630594.1) and *Gryllus bimaculatus* (AB300616.1), selected for the availability of expression data for these homologs. We searched for *Egfr* homologs in the genomes of the spider *Parasteatoda tepidariorum* (GCA_000365465.2), the scorpion *Centruroides sculpturatus* (GCA 000671375.2), the horseshoe crabs *Limulus polyphemus* (GCA_000517525.1) and *Tachypleus tridentatus* (CNA0000821), the mite *Tetranychus urticae* (GCF_000239435.1), the tick *Ixodes scapularis* (GCA_000208615.1), the water-strider *Gerris buenoi* (GCA_001010745.2; OGSv1.0), and the embryonic transcriptome of the pseudoscorpion *Conichochernes crassus*. We additionally included known orthologs of the myriapod *Glomeris marginata* (LT716782) (Janssen, 2017), and used as outgroups vertebrate *Egfr* homologs of *Homo sapiens, Mus musculus, Gallus gallus* and *Danio rerio.* A list of accession numbers is provided in Table S4. The final list of putative *Egfr* homologs was aligned as proteins using Clustal Omega (Sievers et al., 2011) implemented in SeaView v. 5.0.4 (Gouy, Guindon, & Gascuel, 2010), and analyzed under maximum likelihood with IQTREE v. 1.6.12 (-m TEST -bb 1000) (Nguyen, Schmidt, Haeseler, & Minh, 2015). EGFR protein motifs were predicted using the MOTIF tool in GenometNet (https://www.genome.jp/tools/motif) (Kanehisa, 1997).

### Analysis of microRNAs

An initial search for miRNA families in the genome of *P. opilio* and seven other chelicerates (Table S3) used as queries all miRNAs previously reported from the spider *P. tepidariorum,* the tick *Ixodes scapularis* and the mite *Tetranychus urticae* (Leite et al., 2016; Ontano et al., 2020). To recover unique putative harvestmen miRNAs, we conducted a second search using miRNA families not known in spiders, ticks or mites, but which were shared by at least three mandibulate arthropod taxa (insects, centipedes or crustaceans). An initial BLAST search was performed using the following commands: blastn -word_size 4 -reward 2 -penalty −3 – evalue 0.05, with those sequences with e-value < 0.05 and percentage identity > 75% being retained. To further corroborate our selected miRNAs, their structure and minimum free energy were generated with RNAfold v.2.4.13 (part of the ViennaRNA package 2.0 (Lorenz et al., 2011)) using default options. Non-chelicerate miRNAs were obtained from the miRbase (Kozomara, Birgaoanu, & Griffiths-Jones, 2019).

### Cloning, In Situ Hybridization, and Double-Stranded RNA Microinjection

Fragments of *Popi-EgfrA* and *Popi-Dfd* were amplified from cDNA using gene specific primers designed with Primers3 v. 4.1.0 (Koressaar & Remm, 2007) and appended with T7 ends (Fig. S5, S9; Table S5). Amplicon cloning was conducted with TOPO TA Cloning Kit One Shot Top 10 *Escherichia coli* following the manufacturer’s protocol (Invitrogen, CA, USA). All fragments were Sanger sequenced to verify on-target cloning and directionality (Table S6). Double-stranded RNA for *Popi-EgfrA* and for both fragments of *Popi-Dfd* were synthesized using the MEGAScript T7 transcription kit (ThermoFisher, MA, USA). Colorimetric in situ hybridization assays used sense and anti-sense RNA probes labeled with DIG RNA labeling mix (Roche, Basel, Switzerland), following a published procedure (Sharma et al., 2012). Probes for *Po-EgfrA* were synthesized from a PCR template using universal T7 primers and T7 polymerase (New England Biolabs, MA, USA). RNA probes for *Popi-Dfd* were synthetized from the plasmid template of *Popi-Dfd3’* using T7/T3 RNA polymerase (New England Biolabs, MA, USA).

Embryonic RNA interference (RNAi) followed the protocol in (Sharma, Schwager, Giribet, Jockusch, & Extavour, 2013). For all gene knockdown experiments, double-stranded RNA was adjusted to 4 μg/μL and mixed with Rhodamine Dextran B dye (20:1) (Thermo Fisher) for visualization of injected volume. Negative control embryos were injected with distilled water in the same dilution of the dye. For each clutch studied, approximately two-thirds of the embryos were injected with dsRNA and the remaining third with water (control). Four clutches were used for the *Popi-Dfd3’* experiment (n=359), two clutches for *Popi-Dfd5’* (n=189), and two for the *Popi-EgfrA* experiment (n=290). The RNAi experiments for *Popi-Dfd3’* and *Popi-EgfrA* were replicated in at least one extra clutch, which was fixed at a range of embryonic stages to be assayed by in situ hybridization. Maternal RNAi against *Ptep-DfdA* followed the same primer design and experimental procure in Pechmann et al. (Pechmann et al., 2015).

Phenotypes were scored upon hatching. Embryos damaged by assisted hatching or by dehydration were scored as indeterminate and excluded from the analysis (n=148 for *Popi-Dfd* RNAi; n=109 for *Popi-EgfrA*). Given the large proportion of mosaic phenotypes (asymmetric penetrance on different sides of the body), we further refined scoring of the 43 individuals presenting a homeotic phenotype in the *Popi-Dfd3’* experiment by separately scoring each half of the body into two classes: (1) Class II (weak): partial L1-to-pedipalp transformation; L1 tarsus is shorter and undivided, metatarsus is reduced; L2 is mostly normal, sometimes with a shorter tarsus and metatarsus; (2) Class I (strong): complete L1-to-pedipalp transformation; L1 lacks a metatarsus; tarsus is undivided, shorter, thicker and has setation pattern of the pedipalp; L1 mesal surface of the tibia and patella bears dark setal cluster (spurs); L2 is shorter, completely lacks a metatarsus or has a defective metatarsus; tarsomeres are reduced or absent; the podomere is thicker and has pedipalp setation and claw. For *Popi-EgfrA* RNAi, phenotypic effects scored included dorsal body fusion, eye reduction, inter appendage coxal fusion, leg podomere fusion, chelicera finger reduction, pedipalp claw reduction, and leg distal reduction.

## Acknowledgements

RNA sequencing was performed at the Center for Systems Biology, Harvard University. Genomic laboratory work was conducted at the Smithsonian Laboratories of Analytical Biology (LAB). Specimen vouchering and long-term cryogenic curation was provided by the Smithsonian National Museum of Natural History. Microscopy was performed at the Newcomb Imaging Center, Department of Botany, University of Wisconsin-Madison. GG was supported by a Wisconsin Alumni Research Foundation Fall Research Competition award. Computing was conducted by the Smithsonian Institution High Performance Cluster (SI/HPC), Smithsonian Institution (https://doi.org/10.25572/SIHPC). This material is based on work was supported by the Food and Drug Administration (JAC), the Global Genome Initiative Grant No. GGI-Exploratory-2016-047 (VLG), and National Science Foundation grants IOS-1552610 and IOS-2019141 (PPS).

## Author Contributions

Animal husbandry: LBG, EVWS, GG, CMB; Genome sequencing: VLG; RNA sequencing: GG, PPS, EVWS, JB; Genome assembly and annotation: VLG, GG, JB, CESL, PPS; Genome quality assessment: VLG; Orthology inference and phylogenetic analysis: GG, JB; Analysis of microRNAs: CESL; Cloning, in situ hybridization, and double-stranded RNA microinjection: GG, LBG, PPS; Photography: CMB; Designed the study: GG, PPS, VLG; Obtained funding: JAC, PPS, VLG; Drafted the manuscript: GG, PPS, VLG; Revised the manuscript: all authors.

