## Supplementary material for "The genome of a daddy-long-legs (Opiliones) illuminates the evolution of arachnid appendages and chelicerate genome architecture": Fig S; Table S; Supplementary Methods

Guilherme Gainett[1*], Vanessa L. González[2*], Jesús Ballesteros[1], Emily V. W. Setton[1], Caitlin M. Baker[1], Leonardo Barolo Gargiulo [1], Carlos E. Santibáñez-López[3], Jonathan A. Coddington[2], Prashant P. Sharma[1]
[1]Department of Integrative Biology, University of Wisconsin-Madison, Madison, WI, USA 53706
[2] Global Genome Initiative, Smithsonian Institution, National Museum of Natural History, 10th and Constitution, NW Washington, DC 20560-0105, USA
[3] Department of Biology, Eastern Connecticut State University, 83 Windham Street, Willimantic, CT 06226, USA

*co-first authors; corresponding authors

Contains:

Supplementary Methods
Supplementary Figures S1 to S10
Supplementary Tables S1 to S6
Supplementary References

### Supplementary Methods

### Genome Sequencing

Illumina paired end reads were run through kraken v.2.0 (Wood, Lu, & Langmead, 2019) to remove bacterial or viral contaminants prior to use in subsequent analyses:

-run command: kraken2 --db --threads 20 --gzip-compressed --paired

### Genome Assembly

We assembled the long-reads in Canu v1.7 (Koren et al., 2017) using default parameters:

-run command: canu -p phalangium -d phalangium-pacbio useGrid=false ovsMethod=sequential gnuplot=gnuplot gnuplotImageFormat=svg genomesize=1.1g -pacbio-raw phalangium_16cells.fasta

The resulting contigs from Canu were processed using two rounds of scaffolding, gap filling and polishing. Scaffolding was done with SSPACE-LongRead v1.1 (Boetzer & Pirovano, 2014):

-run command: perl SSPACE-LongRead.pl -c scaffolds_canu_kraken_unclassified_pilon.fasta -p phalangium_16cells.fasta -t 40

Followed by gap filling with PBJelly v15.8 (English et al., 2012):

-run command: Jelly.py mapping phalangium_pbjelly.xml; Jelly.py support phalangium_pbjelly.xml; Jelly.py extraction phalangium_pbjelly.xml; Jelly.py assembly -x "--nproc=20" phalangium_pbjelly.xml; Jelly.py output phalangium_pbjelly.xml

And lastly polishing with Pilon v. 1.23 (Walker et al., 2014):

-run command: java -Xmx700G -jar pilon-1.23.jar --genome jelly.out.fasta --frags Illumina_data_kraken_unclassified_mapped_scaffolds_canu_sspacelg1_pbj1_pilon.sorted.bam --output scaffolds_canu_sspacelg2_pbj2_pilon2_kraken_unclassified_pilon --diploid --threads 40

After the second round of polishing the regions arising from haplotypic duplications were identified and removed with Purge_dups v1.2.3 (Guan et al., 2020):

-run command: run_purge_dups.py -p bash phalan_2rounds_pilon.json purge_dups/bin/ scaffolds_canu_sspacelg2_pbj2_pilon2_kraken_unclassified_pilon_purge_dups

### Genome Annotation

Prior to genome annotation, we generated a custom repeat library using RepeatModeler open-1.0.11 (Smit, Hubley, & Green, n.d.):

-run command: RepeatModeler -pa 30 -database Phalangium

Repeats identified were classified using RepeatClassifier open-1.0.11:

-run command: RepeatClassifier -consensi phalangium-families.fa -stockholm phalangium-families.stk -engine ncbi

In order to exclude any protein-coding genes mistakenly classified as repeats, the repeats were compared to known protein coding genes (uniprot) using BLASTx (Altschul, Gish, Miller, Myers, & Lipman, 1990; Camacho et al., 2009). Sequences with hits of e ≤ 0.001 against known protein-coding genes that were also not classified were removed. We used RepeatMasker open-4.0.6 (Smit et al., n.d.) to soft mask repeats and low complexity regions using the custom repeat library generated here combined with known Arachnida repeats:

run command: RepeatMasker -pa 30 -lib phalangium-families.classified.filtered.arachnid_RM_lib_RepeatLibs.fa genome.purged.fa -dir softmask_purge_dups -xsmall -poly

We generated gene predictions using BRAKER2 v.2.1. (Barnett, Garrison, Quinlan, Strömberg, & Marth, 2011; Brůna, Lomsadze, & Borodovsky, 2020; Buchfink, Xie, & Huson, 2015; Gotoh, 2008; Hoff, Lange, Lomsadze, Borodovsky, & Stanke, 2016; Hoff, Lomsadze, Borodovsky, & Stanke, 2019; Iwata & Gotoh, 2012; H. Li et al., 2009; Lomsadze, Burns, & Borodovsky, 2014; Lomsadze, Ter-Hovhannisyan, Chernoff, & Borodovsky, 2005; Stanke, Diekhans, Baertsch, & Haussler, 2008; Stanke, Schöffmann, Morgenstern, & Waack, 2006) with hints generated from previously generated RNAseq reads (NCBI SRX450969) and a curated developmental transcriptome assembly (NCBI PRJNA690950). BRAKER2 is a genome annotation pipeline that allows for automated training of the gene prediction tools GeneMark-EX and AUGUSTUS from RNASeq and protein homology information:

-run command : braker.pl --species=Phalangium_opilio --genome=purged.fa.masked --softmasking --bam=hisat2/purged/Phalangium_assembly_purged_2_noRNAreads.sorted.bam --prot_seq=Popi.longest.pep.txt --cores=32 --GENEMARK_PATH=GENEMARK/gmes_linux_64/ --makehub --email= --etpmode --prg=exonerate).

RNAseq reads were mapped to the masked genome using HISAT2 v.2.2.0 (Kim, Paggi, Park, Bennett, & Salzberg, 2019) using default parameters prior to running BRAKER2:

-run command: hisat2-build purged.fa.masked Phalangium_assembly_purged; hisat2 -x Phalangium_assembly_purged -1 norRNA_1.fq -2 norRNA_2.fq --threads 10 -S Phalangium_assembly_purged_2_noRNAreads.sam --no-unal

Gene predictions were then validated using BLASTx (e ≤ 0.001) against known protein-coding genes (Altschul et al., 1990; Camacho et al., 2009). Functional annotations were identified using Blast2GO (Götz et al., 2008). Following annotation, we removed genes from our annotation that did not generate significant BLAST hits, functional annotation, or lacked transcript evidence. This gene set was then further refined with a 98% similarity threshold using CDHIT (Fu, Niu, Zhu, Wu, & Li, 2012):

-run command: cd-hit -i Blast2Go_annotated_genes_plus_busco.annotated.aa -o Blast2Go_annotated_genes_plus_busco.annotated_CDHIT098.aa -c 0.98 -n 5 -T 4

**Genome Quality Assessment**

Assembly statistic (Table S1) were visualized using Assembly_Stats (Challis, 2017) (Fig. S1). Genome size estimations were conducted in GenomeScope (Vurture et al., 2017) using the generated short read sequence information (results:http://genomescope.org/analysis.php?code=9HRuNixuqcF19W4n6Re6). The final assembly was subjected to contamination screening in Blobtools v.1.0 (Laetsch & Blaxter, 2017) Taxonomic matches, based on sequence similarity search results using BLAST, were categorized according to phylum and were run for both the short read and long read sequence information. This data was used to generate BlobPlots visualizations (Fig. S3). BlobPlots are two-dimensional scatter plots, depicting scaffolds shown as dots and binned by taxonomic match. Each scaffold (dot) is plotted by the base coverage of the sequence by the GC content. Significant spread or clustering in the graph can be used to identify sequences of non-target origin. We used Benchmarking Universal Single-Copy Orthologs (Busco v4.0.2 (Waterhouse et al., 2017)) to assess the assembly completeness by comparing it to 1,013 orthologs contained in the arthropoda_odb10 gene database (Kriventseva et al., 2019).

**Ks plots**

Whole-paranome Ks frequency distributions were calculated with the software *wgd* (Zwaenepoel & Van de Peer, 2019) using average linkage clustering for phylogenetic inference:

-run command: wgd mcl --cds --mcl -s FILENAME -o ./ -n 8 & wgd ksd -o ./ -n 8 FILENAME.mcl FILENAME.cds.fasta

Ks plots generated for the following genomes: *Limulus polyphemus* (GCF_000517525.1) (Battelle et al., 2016), *Tachypleus tridentatus* (CNA0000821) (Gong et al., 2019), *Parasteatoda tepidariorum* (GCF_000365465.2) (Schwager et al., 2017), *Centruroides sculpturatus* (GCF_000671375.1) (Schwager et al., 2017), *Tetranychus urticae*

(GCF_000239435.1) (Grbić et al., 2011), *Galendromus occidentalis* (GCF_000255335.1) (Hoy et al., 2016), *Ixodes scapularis* (GCA_000208615.1) (Gulia-Nuss et al., 2016), *Strigamia maritma* (GCA_000239455.1) (Chipman et al., 2014), and *Phalangium opilio*.

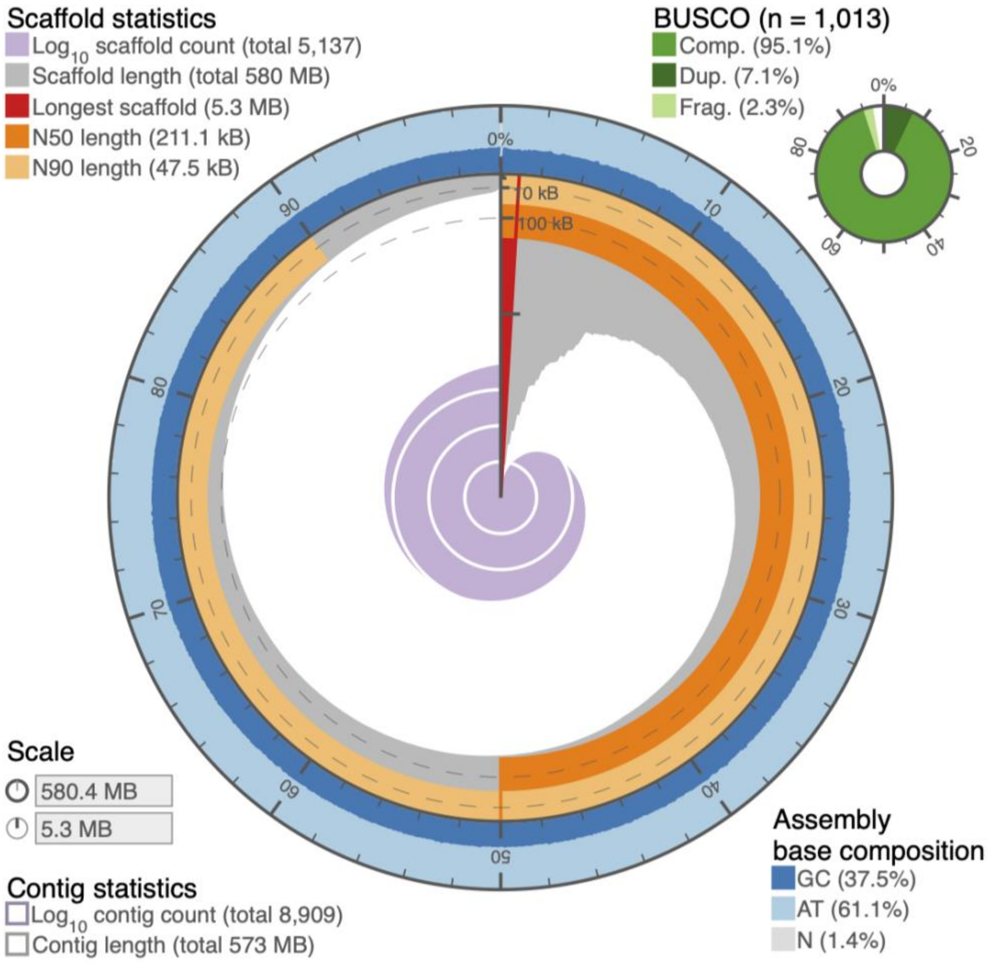


**Fig. S1:** *P. opilio* assembly statistic visualization (Challis, 2017) showing the genome N50 (dark orange), N90 (light orange), base composition (percentage of GC in dark blue, AT in light blue, and N in light grey), and BUSCO results (top right, in shades of green).

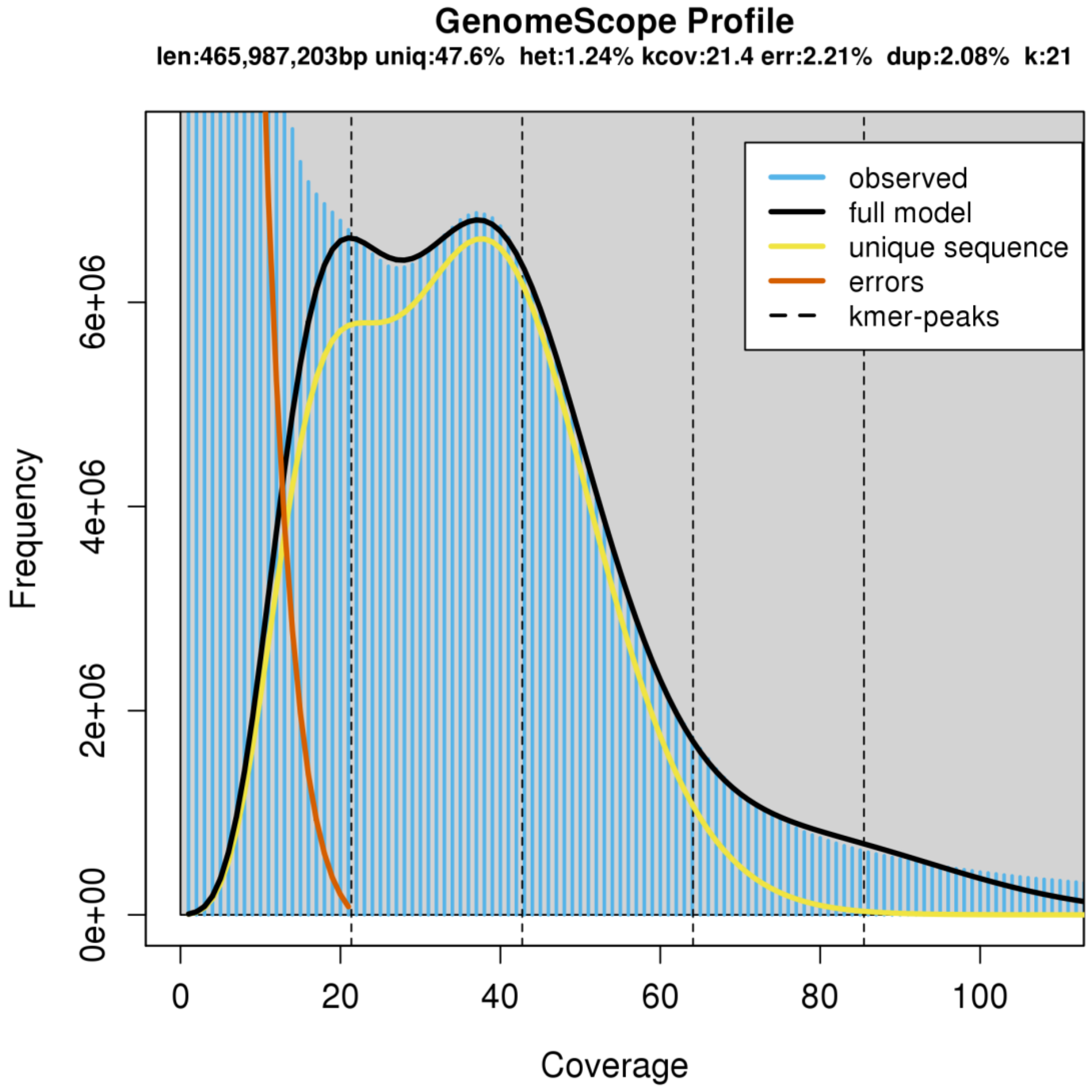


**Fig. S2:** Genomescope (Vurture et al., 2017) plot of the 21-mer k-mer content of the *P. opilio* genome used for genome profiling (genome size, heterozygosity, uniqueness, duplication level) estimated from short read sequence data.

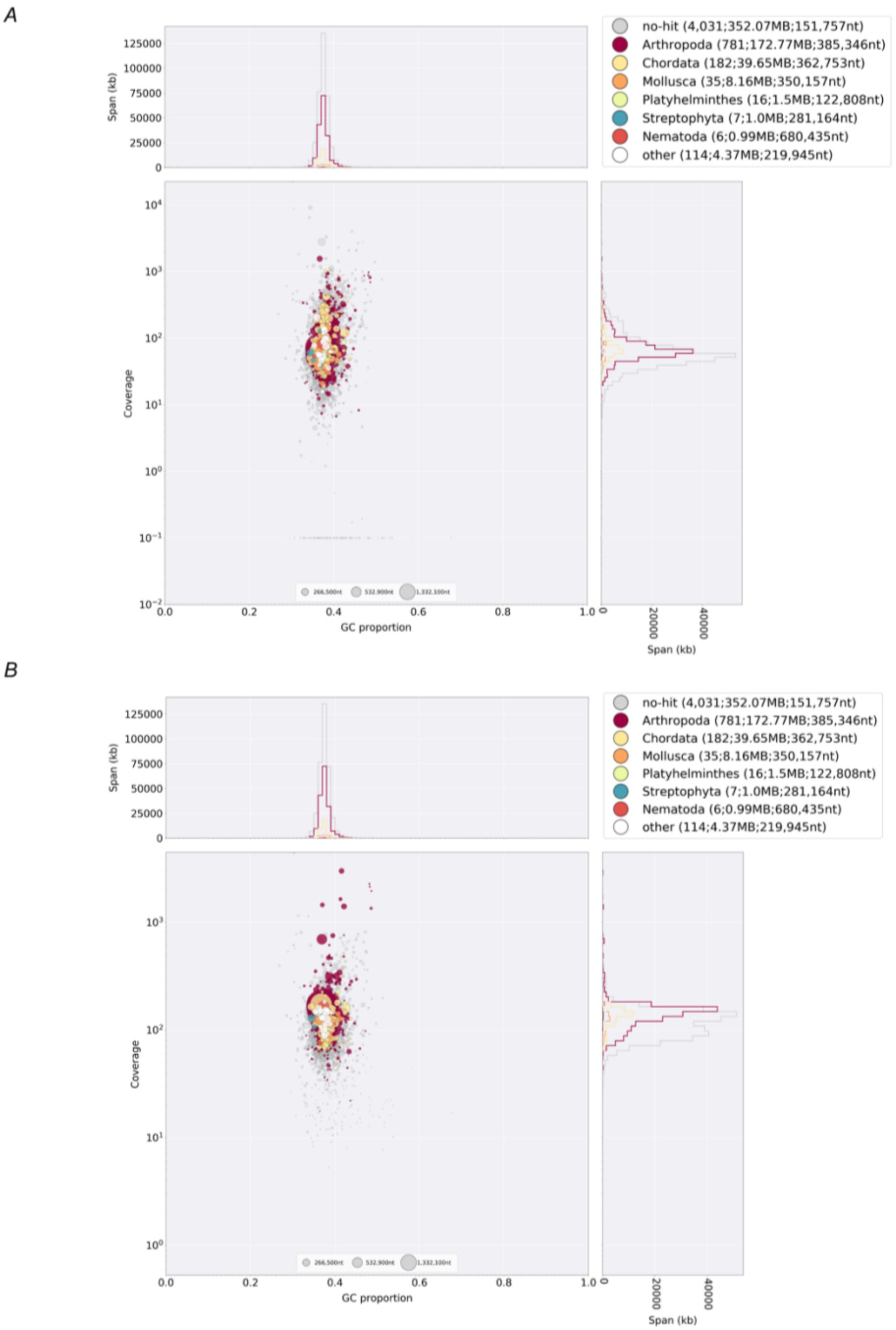


**Fig. S3:** Blobplots (Laetsch & Blaxter, 2017) of the *P. opilio* genome comparing (*A*) coverage calculated based on the short read sequence information and (*B*) coverage shown calculated based on the long read sequence information by the average GC content of each scaffold. Circles are proportional to scaffold length; matches are categorized according to phylum at the taxonomic level.

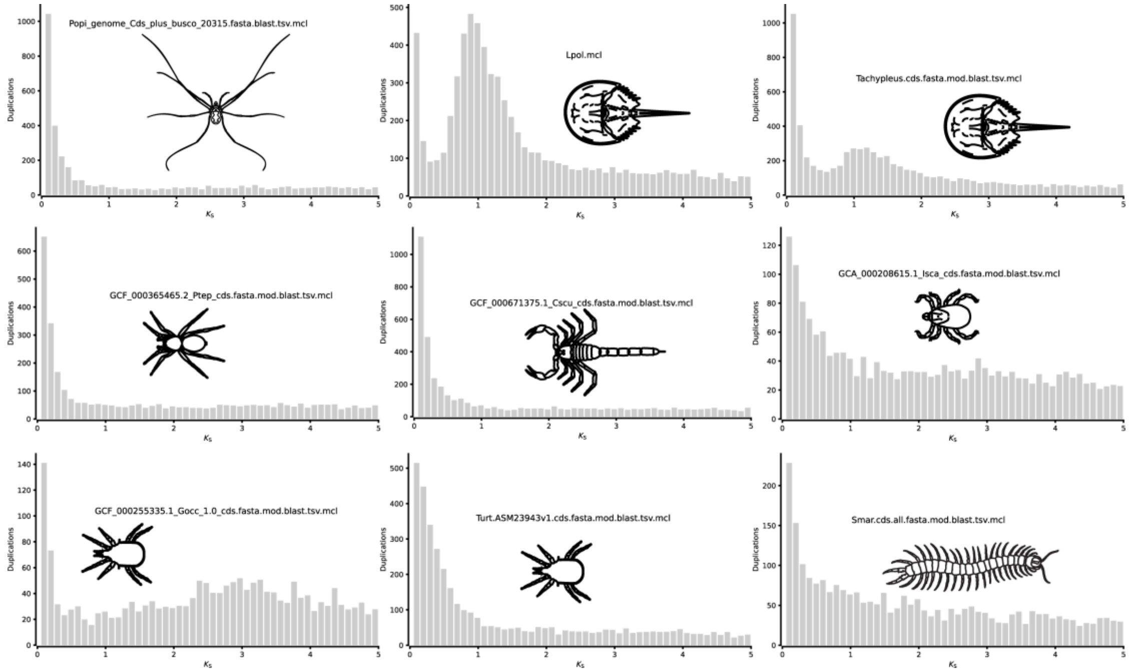


**Fig. S4:** Whole-paranome Ks frequency distributions of *Limulus polyphemus* (*Lpol*), *Parasteatoda tepidariorum* (*Ptep*), *Galendromus occidentalis* (formerly *Metaseiulus*) (*Gocc*), *Tachypleus tridentatus*, *Centruroides sculpturatus* (*Cscu*), *Ixodes scapularis* (*Isca*), *Strigamia maritima* (*Smar*), *Tetranychus urticae* (*Turt*) and *Phalangium opilio* (*Popi*). References to the original genomes are provided in the Supplementary Methods.

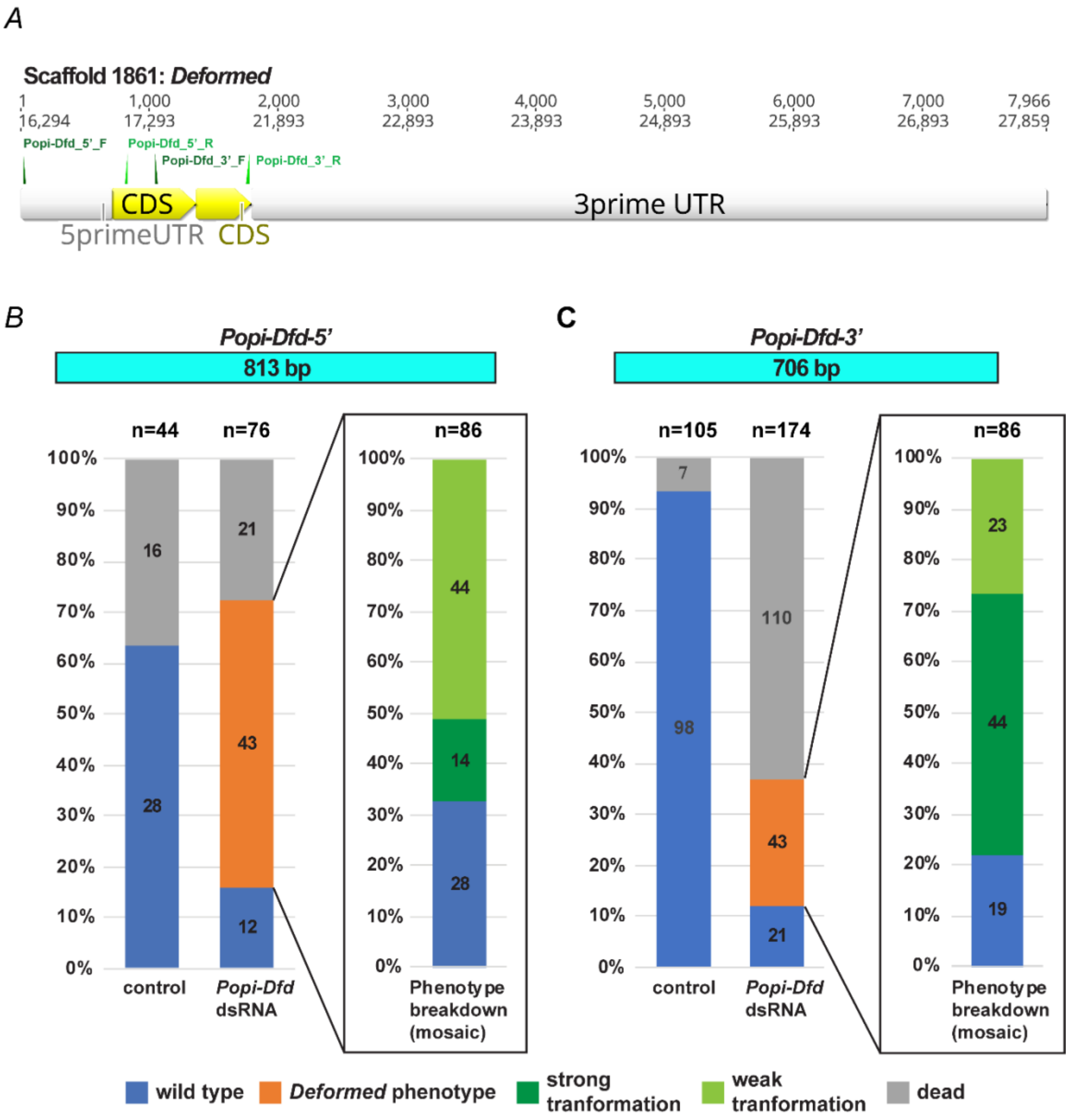


**Fig. S5:** (*A*) Predicted *Po-Deformed* mRNA (Scaffold 1861) and mapped *Po-Dfd 5'* and *Po-Dfd 3'* primers. (*B*) Phenotypic distribution of control and *Po-Dfd* dsRNA RNAi experiments for the non-overlapping fragments.

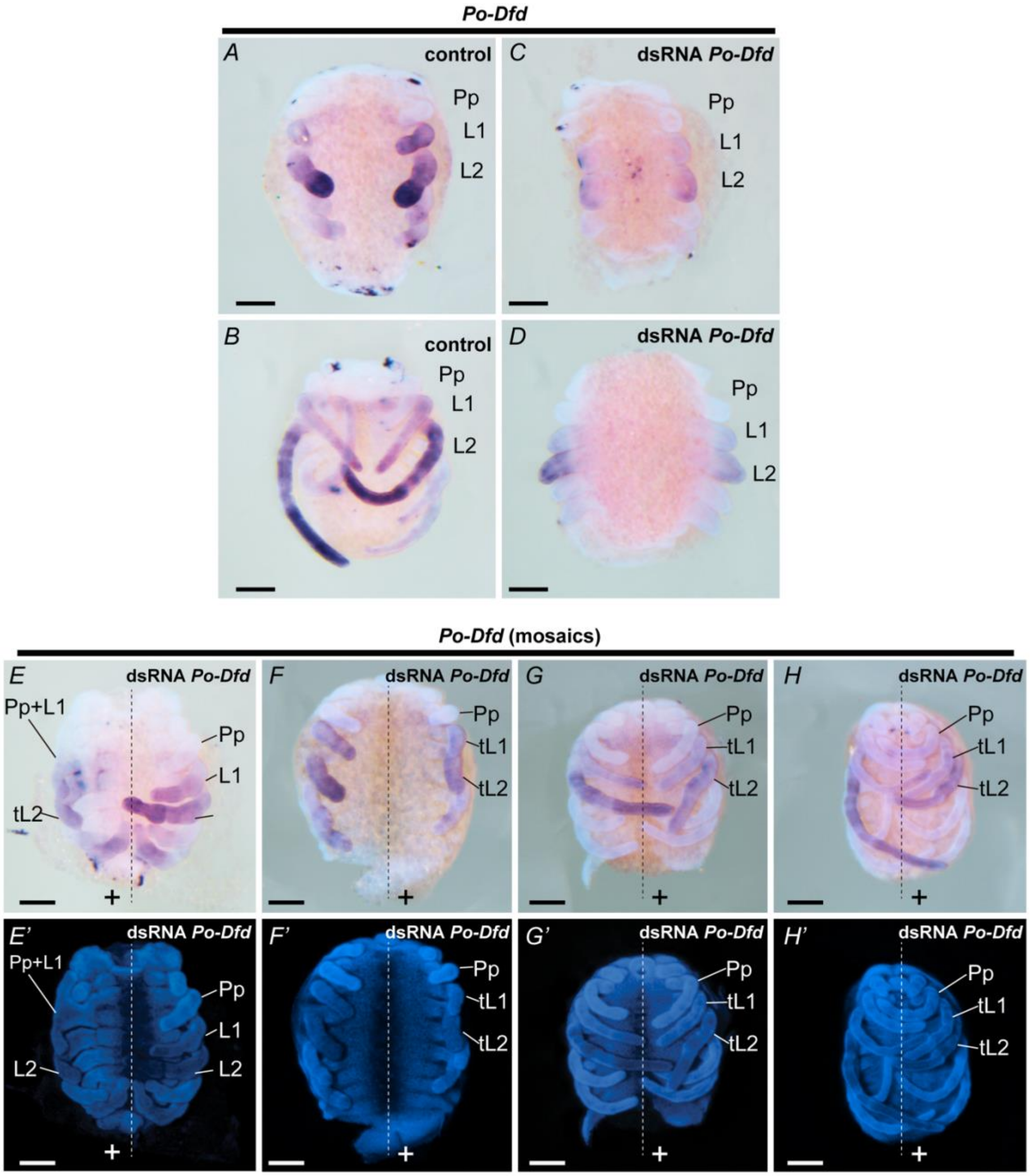


**Fig. S6:** *Po-Dfd* expression (in situ hybridization) in control and *Po-Dfd 3'* dsRNA-injected embryos. Knockdown embryos have decreased expression. (*A* and *B*) Control embryos. (C *and* D) dsRNA *Po-Dfd 3'*-injected embryos. (*E–F*) Mosaic embryos, in which transformed leg 1 and leg 2 (cross sign; shorter appendages than control side) correlate with decreased expression of *Po-Dfd*. L1, leg 1; tL1, transformed L1; L2, leg 2; tL2, transformed leg 2; Pp, pedipalp. Scale bar: 100µm.

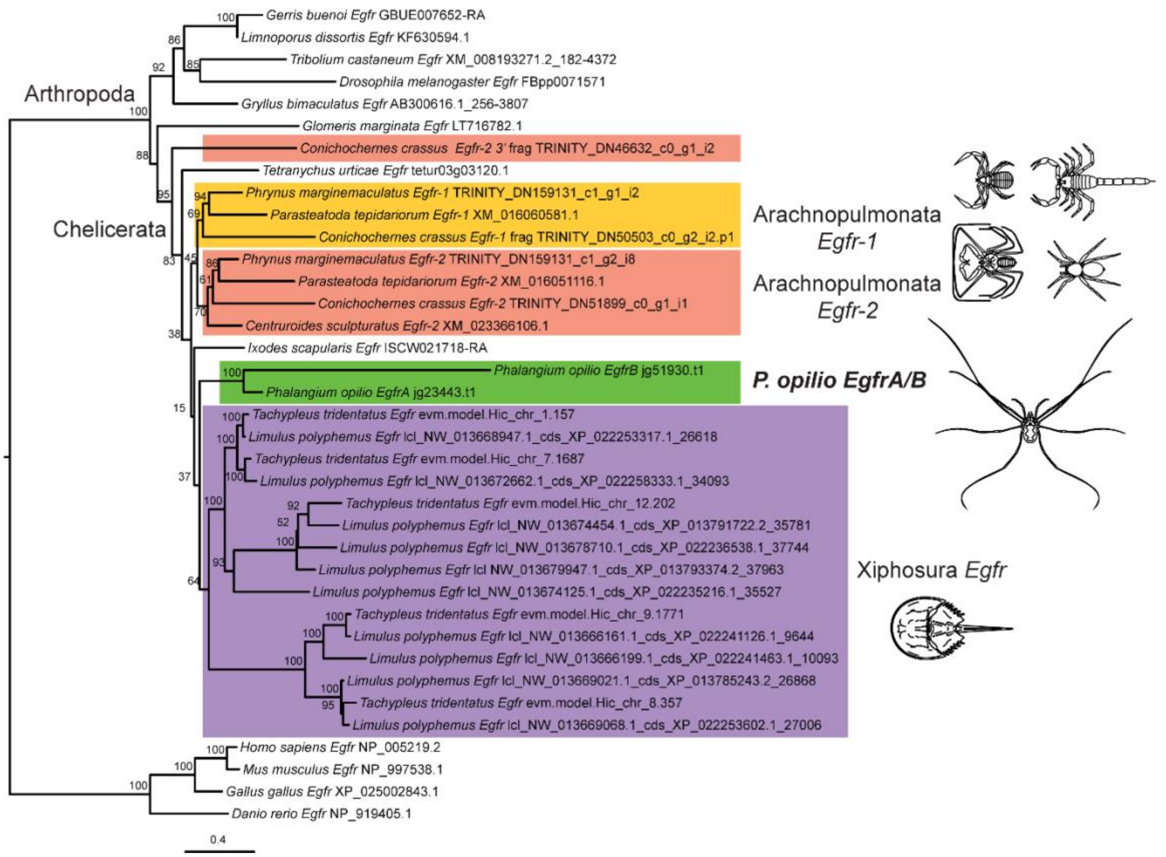


**Fig. S7:** Maximum likelihood tree of *Epidermal growth factor receptor* (*Egfr*) homologs reveals independent duplications in Chelicerata. Values on the nodes are ultrafast bootstraps. The accession numbers to the original sequences are provided in Table S4.

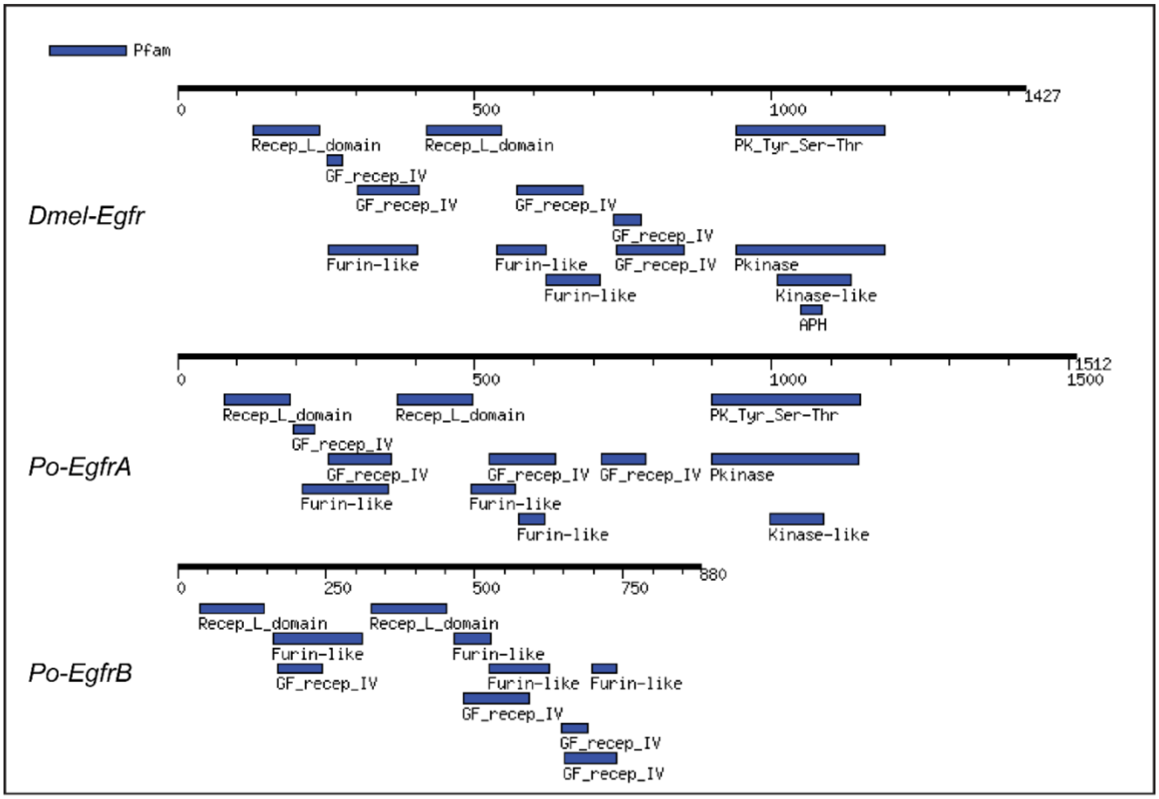


**Fig. S8:** Protein motif predictions for *Dmel-Egfr* (*Drosophila melanogaster*), *Po-EgfrA*, and *Po-EgfrB* (*P. opilio*). Generated with MOTIF tool in GenometNet (https://www.genome.jp/tools/motif). PK_Tyr_Ser-Thr, Protein tyrosine and serine/threonine kinase (PF07714); Recep_L_domain, Receptor L domain (PF01030); GF_recep_IV, Growth factor receptor domain IV (PF14843); Pkinase, Protein kinase domain (PF00069); Furin-like, Furin-like cysteine rich region (PF00757); Kinase-like, Kinase-like (PF14531).

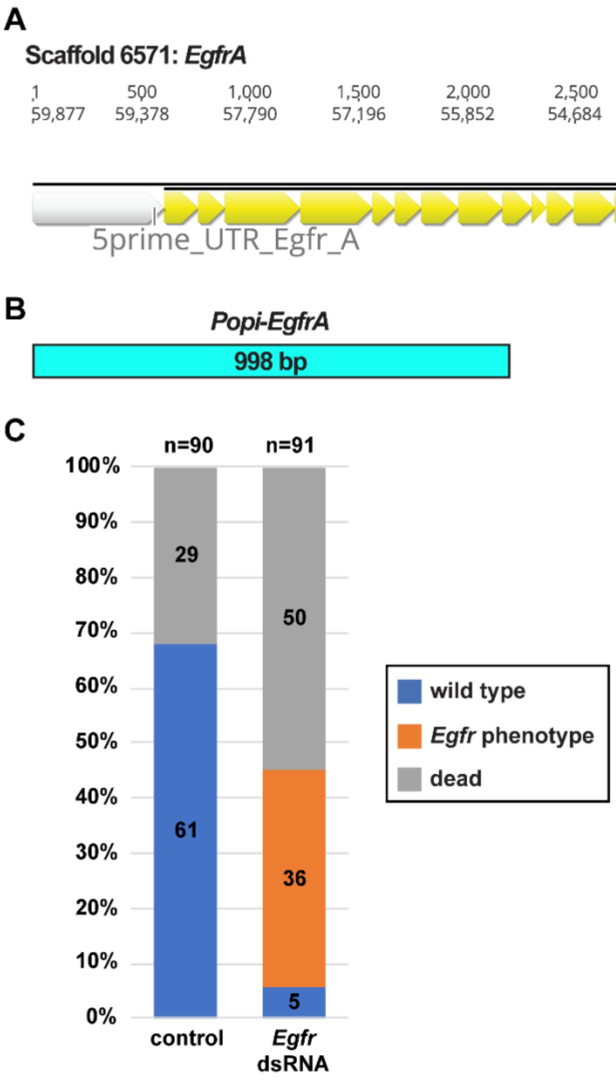


**Fig. S9:** (*A*) Predicted *Po-EgfrA* mRNA (Scaffold 6571) and mapped *Po-EgfrA* primers. (*B*) Phenotypic distribution of control and *Po-Egfr* dsRNA RNAi experiment.

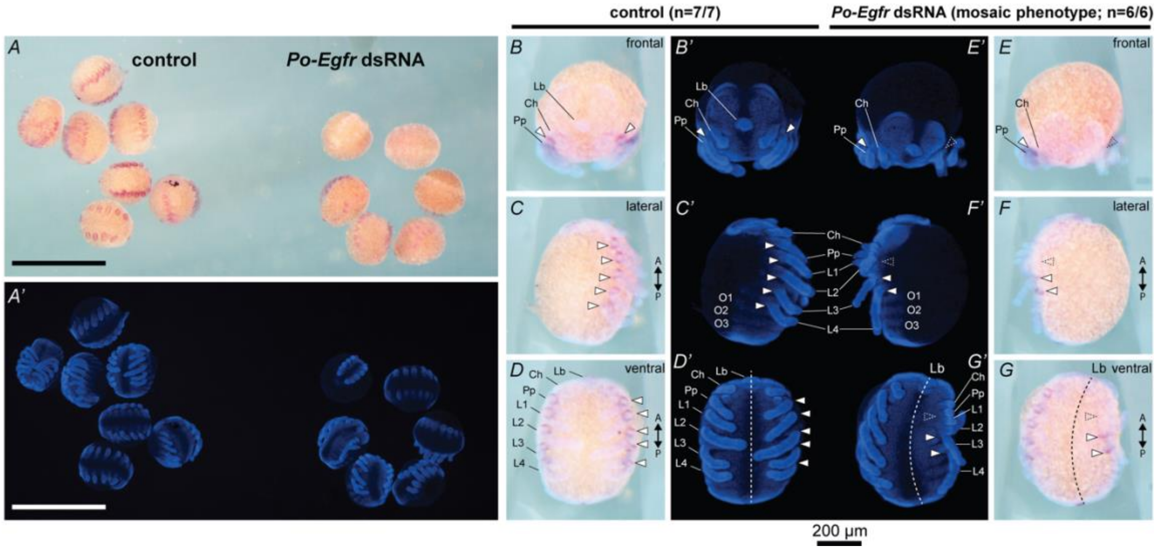


**Fig. S10:** *Po-EgfrA* expression (in situ hybridization) in control and *Po-EgfrA* dsRNA-injected embryos. Knockdown embryos have decreased expression. (*A*) Control (left) and dsRNA-injected (right) embryos. (*B*–*D*) Control embryo in frontal *(B*), lateral (*C*), and ventral (*D*) view. (*E*–*G*) dsRNA-injected embryos in frontal *(E*), lateral (*F*), and ventral (*G*) view. Scale bar: 1 mm (*A* and *A'*); 200 µm (*B*–*G*).

**Table S1:** Assembly statistics for the *P. opilio* genome.

| Contig statistics | Assembly (Prior to Purge_dups) | Post Purge_Dups |
|---|---|---|
| Total number of base pairs | 869216701 | 572607079 |
| Total number of contigs | 17856 | **8349** |
| N10 | 411768 | 570744 |
| N20 | 246790 | 314688 |
| N30 | 172934 | 220382 |
| N40 | 127429 | 165420 |
| N50 | 91196 | **127429** |
| L10 | 106 | 46 |
| L20 | 388 | 189 |
| L30 | 812 | 410 |
| L40 | 1400 | 711 |
| L50 | 2208 | 1106 |
| GC content | 37.58% | 37.51% |
| Mean scaffold size | 24947.5 | 35724 |
| Median scaffold size | 48679.25 | 68584 |
| Longest contig is | 4894628 | 4876499 |
| **Scaffold statistics** | | |
| Total number of base pairs | 881679069 | 580443427 |
| Total number of scaffolds | 9595 | **5137** |
| N10 | 658428 | 790435 |
| N20 | 435595 | 481317 |
| N30 | 327795 | 352528 |
| N40 | 248444 | 269097 |
| N50 | 191854 | **211089** |
| L10 | 73 | 36 |
| L20 | 239 | 135 |
| L30 | 472 | 279 |
| L40 | 784 | 470 |
| L50 | 1191 | 715 |
| GC content | 37.05% | 37.50% |
| Mean scaffold size | 43557 | 112992.7 |
| Median scaffold size | 91889.43 | 58814 |
| Longest scaffold is | 5368172 | 5328218 |
| **BUSCO (arthropoda_ob10)** | | |
| Complete BUSCOs (C) | 96.5% (978) | 95.1% (963) |
| Complete and single-copy BUSCOs (S) | 44.1% (447) | **88.0% (891)** |
| Complete and duplicated BUSCOs (D) | 52.4% (531) | **7.1% (72)** |
| Fragmented BUSCOs (F) | 1.6% (16) | 2.3% (23) |
| Missing BUSCOs (M) | 1.9% (19) | 2.6% (27) |
| Total BUSCO groups searched | 1013 | 1013 |

**Table S2:** List of manually annotated genes from *P. opilio* genome.

| Gene | BREAKER annotation | Genome scaffold | Transcriptome 1 | Transcriptome 2 | Comments |
|---|---|---|---|---|---|
| **Hox genes** | | | | | |
| *labial* | jg1548.t1 | Contig9232_pilon | comp42325_c3_seq5 | comp184606_c0_seq1<br>comp155184_c0_seq1 | |
| *proboscipedia* | jg1352.t1 | Contig3674_pilon | comp41843_c1_seq3<br>comp41843_c0_seq4 | comp92303_c0_seq2<br>comp157201_c0_seq2<br>comp132082_c0_seq1<br>comp165514_c0_seq1 | |
| *Hox3* | jg35409.t1 | Contig4435_pilon | comp40437_c0_seq6<br>comp40437_c0_seq2<br>comp40437_c0_seq1 | comp163899_c0_seq1<br>comp187324_c1_seq8<br>comp187324_c1_seq6<br>comp187324_c1_seq3 | |
| *Deformed* | jg40596.t1 | Contig1861_pilon | comp41255_c0_seq1 | comp185125_c0_seq9<br>comp181322_c1_seq1 | |
| *Sex combs reduced* | jg14111.t1 | Contig8598_pilon | comp44161_c2_seq6<br>comp44161_c2_seq2 | comp175128_c0 | |
| *fushi tarazu* | jg14110.t1 | Contig8598_pilon | comp33981_c0_seq2<br>comp33981_c0_seq4<br>comp33981_c0_seq3 | comp181615_c1_seq1<br>comp142510_c0_seq2<br>comp173869_c1_seq3<br>comp173869_c1_seq2 | |
| *Antennapedia* | jg14106.t1 | Contig8598_pilon | comp40182_c0_seq3<br>comp40182_c0_seq1 | comp171674_c0_seq1<br>comp171674_c0_seq2<br>comp178701_c0_seq1<br>comp178701_c0_seq2 | |
| *Ultrabithorax* | jg14096.t1 | Contig8598_pilon | comp37908_c1_seq1<br>comp37908_c1_seq2<br>comp37908_c0_seq1 | comp187200_c1_seq2<br>comp182847_c1_seq6<br>comp182847_c1_seq8 | |
| *abdominal-A* | jg14091.t1 | Contig8598_pilon | comp31172_c0_seq1 | comp170823_c0_seq2 | |
| | | | comp31172_c0_seq2 | comp170823_c0_seq1 | |
| *Abdominal-B* | jg14089.t1 | Contig8598_pilon | comp38644_c0_seq5<br>comp38644_c0_seq4<br>comp38644_c0_seq3<br>comp38644_c0_seq2<br>comp38644_c0_seq1 | comp176401_c0_seq3<br>comp176401_c0_seq4<br>comp176401_c0_seq5<br>comp176401_c0_seq2<br>comp176401_c0_seq1 | |
| **EGFR** | | | | | |
| *Egfr-A* | jg23443.t1 | Contig6571_pilon | comp44061_c1_seq1<br>comp44061_c1_seq3<br>comp44061_c0_seq3<br>comp44061_c1_seq4<br>comp44061_c1_seq5; | comp175413_c0_seq1<br>comp175413_c0_seq2<br>comp188269_c0_seq1<br>comp188269_c0_seq2 | |
| *Egfr-B* | jg51930.t1 | Contig5000_pilon | comp33899_c0_seq1; | comp177011_c0_seq1<br>comp177011_c0_seq4<br>comp177011_c0_seq7<br>comp177011_c0_seq8<br>comp177011_c0_seq2<br>comp177011_c0_seq6 | |

| | | | | | |
|---|---|---|---|---|---|
| **Leg gap genes** | | | | | |
| *homothorax* | not predicted | Contig890_pilon | | comp175377_c0_seq8 | partial, 5' fragment |
| | not predicted | Contig5128_pilon | | | partial |
| | not predicted | Contig1882_pilon | | | partial |
| | not predicted | Contig5036_pilon | | | partial, 3' fragment |
| *dachshund* | not predicted | Contig3437_pilon | comp32064_c0_seq1 | | |
| | | | comp37387_c1_seq2 | | |
| *extradenticle* | not predicted | Contig5455_pilon | | | |
| *spineless* | not predicted | Contig8719_pilon | | | |
| **RDGN genes** | | | | | |
| *sineoculis* | jg43040.t1 | Contig598_pilon | comp39452_c0_seq1 | comp183481_c0_seq1 | 3' fragment |
| | jg130.t1 | Contig2734_pilon | comp39452_c0_seq3 | comp183481_c0_seq1 | 5' fragment |
| | | | comp39452_c0_seq4 | comp183481_c0_seq10 | |
| | | | | comp183481_c0_seq10 | |
| | | | | comp183481_c0_seq11 | |
| | | | | comp183481_c0_seq11 | |
| | | | | comp183481_c0_seq12 | |
| | | | | comp183481_c0_seq13 | |
| | | | | comp183481_c0_seq13 | |
| | | | | comp183481_c0_seq2 | |
| | | | | comp183481_c0_seq2 | |
| | | | | comp183481_c0_seq3 | |
| | | | | comp183481_c0_seq4 | |
| | | | | comp183481_c0_seq5 | |
| | | | | comp183481_c0_seq7 | |
| | | | | comp183481_c0_seq7 | |
| | | | | comp183481_c0_seq8 | |
| | | | | comp183481_c0_seq9 | |
| *Optix* | jg30497.t1 | Contig863_pilon | comp42503_c0_seq3 | comp165591_c0_seq2 | |
| | | | comp42503_c1_seq14 | | |
| *orthodenticle* | jg181.t1 | Contig7625_pilon | comp42066_c0_seq1 | comp185461_c0_seq13 | |
| | | | | comp185461_c0_seq3 | |
| | | | | comp185461_c0_seq6 | |

<insert page break here>

**Table S3:** List of species sampled in the *miRNA* genome survey with accession numbers. Species in bold have not been previously analysed in (Leite et al., 2016; Ontano et al., 2020).

| Order | Species | NCBI accession |
|---|---|---|
| Araneae | ***Araneus ventricosus*** | GCA_013235015.1 |
| | *Acanthoscurria geniculata* | GCA_000661875.1 |
| | ***Stegodyphus dumicola*** | GCA_010614865.1 |
| | *Stegodyphus mimosarum* | GCA_000611955.2 |
| | *Parasteatoda tepidariorum* | GCA_000365465.3 |
| | ***Dysdera silvatica*** | GCA_006491805.1 |
| Scorpiones | *Centruroides sculpturatus* | GCF_000671375.1 |
| | *Mesobuthus martensii* | GCA_000484575.1 |
| Pseudoscorpiones | *Cordylochernes scorpioides* | GCA_003123905.1 |
| Xiphosura | *Carcinoscorpius rotundicauda* | GCA_011833715.1 |
| | *Limulus polyphemus* | GCA_000517525.1 |
| | ***Tachypleus tridentatus*** | GCA_004210375.1 |
| | ***Tachypleus gigas*** | GCA_014155125.1 |
| Acariformes | ***Dinothrombium tinctorium*** | GCA_003675995.1 |
| | ***Brevipalpus yothersi*** | GCA_003956705.1 |
| | *Tetranychus urticae* | GCA_000239435.1 |
| Parasitiformes | *Galendromus occidentalis* | GCA_000255335.1 |
| | *Rhipicephalus microplus* | GCA_013339725.1 |
| | *Varroa destructor* | GCA_002443255.1 |
| | *Ixodes scapularis* | GCA_002892825.2 |

**Table S4:** List of terminals used in the *Egfr* phylogenetic analysis with accession numbers.

| Species | Gene | ID | Database |
|---|---|---|---|
| *Centruroides sculpturatus* | *Egfr* | XM_023366106.1 | NCBI; GCF_000671375.1 |
| *Conichochernes crassus* | *Egfr-1* | TRINITY_DN50503_c0_g2_i2 | NCBI; SRR13003861-2 |
| | *Egfr-2 5' frag* | TRINITY_DN51899_c0_g1_i1 | |
| | *Egfr-2 3' frag* | TRINITY_DN46632_c0_g1_i2 | |
| *Danio rerio* | *Egfr* | NP_919405.1 | NCBI |
| *Drosophila melanogaster* | *Egfr* | FBpp0071571 | Flybase |
| *Gallus gallus* | *Egfr* | XP_025002843.1 | NCBI |
| *Gerris buenoi* | *Egfr* | GBUE007652-RA-CDS | NCBI |
| *Glomeris marginata* | *Egfr* | LT716782.1 | NCBI |
| *Gryllus bimaculatus* | *Egfr* | AB300616.1 | NCBI |
| *Homo sapiens* | *Egfr* | NP_005219.2 | NCBI |
| *Ixodes scapularis* | *Egfr* | ISCW021718-RA | NCBI; GCA_000208615.1 |
| *Limnoporus dissortis* | *Egfr* | KF630594.1 | NCBI |
| *Limulus polyphemus* | *Egfr* | XP_022253602.1 | NCBI; GCF_000517525.1 |
| | | XP_013785243.2 | |
| | | XP_022241126.1 | |
| | | XP_022241463.1 | |
| | | XP_013793374.2 | |
| | | XP_022253317.1 | |
| | | XP_022258333.1 | |
| | | XP_022235216.1 | |
| | | XP_013791722.2 | |
| | | XP_022236538.1 | |
| *Mus musculus* | *Egfr* | NP_997538.1 | |
| *Parasteatoda tepidariorum* | *Egfr-1* | XM_016060581.1 | NCBI; GCF_000365465.2 |
| *Parasteatoda tepidariorum* | *Egfr-2* | XM_016051116.1 | |
| *Phrynus marginemaculatus* | *Egfr-1* | TRINITY_DN159131_c1_g1_i2 | NCBI; SRR12232018 |
| *Phrynus marginemaculatus* | *Egfr-2* | TRINITY_DN159131_c1_g2_i8 | |
| *Tachypleus tridentatus* | *Egfr* | evm.model.Hic_chr_8.357 | CNSA; CNA0000821 |
| | | evm.model.Hic_chr_9.1771 | |
| | | evm.model.Hic_chr_1.157 | |
| | | evm.model.Hic_chr_7.1687 | |
| | | evm.model.Hic_chr_12.202 | |
| *Tetranychus urticae* | *Egfr* | XM_015925527.2; tetur03g03120.1 | NCBI; GCF_000239435.1 |
| *Tribolium casteaneum* | *Egfr* | XM_008193271.2 | NCBI |

**Table S5:** List of primers and amplicons.

| Gene | Primer ID | Forward | Reverse | Annealing temperature (ºC) | Cycle length (s) | Product size (bp) |
|---|---|---|---|---|---|---|
| *Po-Deformed* | Popi_Dfd_T7 (3') | ggccgcggTGTAGTCTGACCTCTTCCGC | cccggggcCGTATTCGGTTTTGGGCTCC | 57 | 30 | 706 |
| *Po-Deformed* | Popi_Dfd_5' | ggccgcggTTTCTGCCGCTACGACTTTG | cccggggcTACGGCGGAGAGTTCATCAA | 57 | 30 | 812 |
| *Po-EgfrA* | Popi_Egfr (A) | ggccgcggGTAGAGGATGTTACGGCCCA | cccggggcCACAGTAACCCCAAAAGCCC | 57 | 30 | 998 |
| *Po-EgfrB* | Popi_Egfr_B_911bp | ggccgcggATGCTCAAAGTGCGACGATC | cccggggcGACCTTGAACCTGTTGCTCG | 57 | 30 | 911 |

**Table S6:** List of cloned sequences used in the RNAi experiments.

| Gene | Primer | Plasmid # | Plasmid ID | Direction | Sequence |
|---|---|---|---|---|---|
| *Po-Deformed* | Popi_Dfd_T7 (3’) | 9 | AXW386 | Forward | NNNNNNNNNNNGGGCGANTGANTTTAGCGGCCGCGAATTCGCCCTTCGGTGTAGTCTGACCTCTTCCGCGGGACCACCGATTCCCAGAGGTTCCGCCGG<br>ACTACCGCCGTCCGTCGACCATGTCGCGAATAGTCACAGTCCCGTTCCGGGTTTGCACGGACCTTTGCCTCAGCACCACGCGCATAGGTCACCCGTTTCGTC<br>ACCGCCACCCAACGGTGGACTGCCAAGGTCCCAACCACCCCAAACGACGTCGTCGCATCCGCCGCAACAGGGCAGTCCGGGCTTACCCAACCCCGATTGCG<br>CCCTTTCGTCCGGCGGACAGCCAGTCATATACCCCTGGATGAAAAAAGTCCATATTAATGCAGCGAACGGAGCGTTCGCGGGGATGGAACCGAAACGCCA<br>ACGGACGGCGTACACGAGGCATCAGATTCTCGAGCTAGAAAAAGAGTTCCATTTTAACCGGTACCTGACTAGAAGACGACGGATTGAAATCGCTCACGCGC<br>TCTGTTTATCCGAGAGGCAAATTAAGATTTGGTTCCAAAATAGGCGAATGAAATGGAAAAAGGACAATAAATTGCCTAATACCAAAAACGTCAAGAAAAAC<br>GCTCAGAAGCAAGCGCAACAACAACAAGCGCAACAGCAACACGCGCAAAATGCCATTAACCATCAGGACATGAGCGGTTTGGCTCAAGCCTCGCCCGCGC<br>AGGGACACGGTCTCTCCCACGCCCCTCAAATGGAGCCCAAAACNAA |
| *Po-Deformed* | Popi_Dfd_5' | 1 | AXW413 | Reverse | NNNNNNNNNNNNNNNNNNTNANCNNNNNTNNNGGACTAGTCCTGCAGGTTTAAACGAATTCGCCCTTCCCGGGGCTACGNCGNNNNGTTCATCAAAAA<br>CGAACTCATGATCATTAATTTTTGGGTGAATTGGATGGAAATAATTTTTCTTCGGGGTTGTCGTTTAACCGCGCGCTGGCTCCGCCTCTCGGCAGAAATTAGC<br>ATATCACGCGGCCGCCGTCGTCCAATCGGCTCGGTGGGCTTTGACCCGGGGCGCTCGCGTCAGATTGATCGATGACAGTGACACATTGCGAGAAAATGGC<br>GGCGCCGGCCATACCTGTAGGCTAGGCCCCGCCGCCCCGCCAACGGACGCGTGAAAATATATTTTTCAAACACAATATACTCGAAAACGCGCTCGTCGCGC<br>GCTCTAGGATCTCGCGCCGCCGCCGAAATTCACCACAACAGTTCCTCGTGCAAAATCCCCGTTCTTGAGTGGAAATGCGATGCAAAAATGATGCCTACCGTA<br>ATGTCCTCAGGCTCGAACACCAACAATCAAGCGTGCAAAATGCTTTATCAGCCGATGTACTCGGTCCTTAAGGGAGAACTCGGGAAGACTCCGAGCGATTG<br>GAACAACTACGGAGATTGTTGTTGGGTCGAGCTAAATTGAAAGCTCTTTCGTTTCGATTAAAAAAAAGGAGCCCGTTGAAGAAGGGCTCGGACGAGTCGTT<br>AAGAGCGCTTGATACGACTCGTGGTGGAGGTTTGAAACGGCAGCCGCGAAGTGCAACTTCTAACGTGGATATGACGTTAAGAGCCGANTTTCGCGCTAAC<br>CCACACCAACTATGTCTGATCAACGTGTGCCAGTAGCAACGTGCACAAAGTCGTAGCGGGCAGAAACCGCGGCCAAGGGCGATTNNGCGGCCGCTAAATT<br>NNAATTCGCCCNTATAGTGAGTCGTATTNNCATTCNNCTGGGCCGTCGTTTTANACGTCNNGACNNGGNNAAANNTGGNNGTTACNNACTTANNCNCNN<br>GCAGCCACATTCCCCTTNNGCNNNCNNGGNNNNATNNCNANAGGNCNNNNNCGANNNCCCNNNNCNAACNNTTNNNNNNCNNNNNCNNNNCNGN<br>ANTNNNNNNNNCNNNNNNNNNNNANNTNNNNNNNNTNNNNNNNNNNTTGNNGNNNNGTNNNNNNNNNGGNANNNNTNN |
| *Po-Deformed* | Popi_Dfd_5' | 2 | AXW414 | Reverse | NNNNNNNNNNNNNNNNNNNANNNNCACTNNNGGACTAGTCCTGCAGGTTTAAACGAATTCGCCCTTCCCGGGGCTANNNCNNANAGTTCATCAAAAAC<br>GAACTCATGATCATTAATTTTTGGGTGAATTGGATGGAAATAATTTTTCTTCGGGGTTGTCGTTTAACCGCGCGCTGGCTCCGCCTCTCGGCAGAAATTAGCA<br>TATCACGCGGCCGCCGTCGTCCAATCGGCTCGGTGGGCTTTGACCCGGAGCGCTCGCGTCAGATTGATCGATGACAGTGACACATTGCGAGAAAATGGCG<br>GCGCCGGCCATACCTGTAGGCTAGGCCCCGCCGCCCCGCCAACGGACGCGTGAAAATATATTTTTCAAACACAATATACTCGAAAACGCGCTCGTCGCGCG<br>CTCTAGGATCTCGCGCCGCTGCCGAAATTCACCACAACAGTTCCTCGTGCAAAATCCCCGTTCTTGAGTGGAAATGCGATGCAAAAATGATGCCTACCGTAA<br>TGTCCTCAGGCTCGAACACCAACAAACGAGCGTGCAAAATGCTTTATCAGCCGATGTACTCGGTCCTTAAGGGAGAACTCCGAGCGATTGGAGCAACTACG<br>GAGATTGTTGTTGGGTCGAGCTAAATTGAAAGCTCTTTCGTTTCGATTAAAAAAGGAGCCCGTTGAGGAAGGGCTCGGACGAGTCGTTAAGAGCGCTTGAT<br>GCCGTTCAACGTTGCTATACGACTCGTGGTGGAGGTTTGAAACGGCAGCCGCGAAGTGCAACTTCTAACGTGGATATGACGTTAAGAGCCGATTTCGCGCT<br>AACCCACACCAACTATGTCTGATCAACGTGTGCCAGTAGCAACGTGCACAAAGTCGTAGCGGCAGAAACCGCGGCCAAGGGCGAATTCGCGGCCGCTAAA<br>TTCAATTCGCCCTATAGTGAGTCGTATTACAATTCACTGGCCGTCGTTTTACAACGTCGTGACTGGGAAAACCCTGGNGTTACCCAACTTAATCGCCTTGCAG<br>CNNATCCCCCTTTCGCCAGCTNNNNNTANNANCNAAGAGGCCCGCACCGATCGCCCTTCCCAACAGTTGGCCAGCCTNNACGTACGGNNGGTTTAAGGTT<br>TANCCNNAAAAANGAAAGCCGTTNNNNNNGGNTTT |
| *Po-EgfrA* | Popi_Egfr | 1 | AXW565 | Forward | NNNNNNNNNNGCGATTGANTTAGCGGCCGCGAATTCGCCCTTGGCCGCGGGTAGAGGATGTTACGGCCCAACAACATCACATTGTCACGCCTGTAGAAA<br>TTTTAGAGTTTACTTCGCTAACAATCAAACAGCTTTTAATTGCACTGCATCATGCCCGGAAGAGATGCCTTACAAAAAATTCGATGACGAGAAAGGCGATCC<br>CTACTGTTCAACTGATGATCCAAATGCTGTTGCTGTACATGGAGCTGAAGATGACCATGTACCTGCCATTTTGGGTGGCACTATTGGTTGTGTGACGCTTTTG<br>GTTATTTTCTTTGCGGTGATGAGCTACCAGTGGGTGCAAAGAGCGAAAACAAAGGAAAACACCATGAAGATGACCATGCGTATGAGTGGATTTGAAGACG<br>AACCATTAAAGCCAAGTAATATCAAACCAAATCTGGCAAAGCTGAGAATTGTAAAGGAACAAGAACTAAGGAAAGGTGGAATCCTTGGTTATGGTGCTTTT<br>GGAACTGTATACCAAGGTGTGTGGCTGCCTGAAGGAGATAATGTGAAAATACCTGTTGCTATTAAAGTTCTTCGTGAAGGAACCTGTCCAAGTACAAATAA<br>AGATTTCCTTGAAGAGGCCTATATAATGGCTAGTGTTGATCATCCAAATGTTCTGAAGCTGTTGGCAGTTAGCTTGGCATCGCAACTCATGTTAGTCACGCA<br>GTTAATGCCTCTGGGCTGCCTTTTAGACTATGTTCGCAATAACCAAGATAAAATTGGACCCAAGCCACTACTAAATTGGTGCACACAGATTGCGAAGGGGAA<br>TGGCTTACTTGGAAGAAAAAAGAATGGTTCATCGTNGATTTGGGCTCTTCGNAATGNGGTTACTGC |
